# Discovering candidate imprinted genes and Imprinting Control Regions in the human genome

**DOI:** 10.1101/678151

**Authors:** Minou Bina

## Abstract

Genomic imprinting is a process thereby a subset of genes is expressed in a parent-of-origin specific manner. This evolutionary novelty is restricted to mammals and controlled by genomic DNA segments known as Imprinting Control Regions (ICRs). The known imprinted genes function in many important developmental and postnatal processes including organogenesis, neurogenesis, and fertility. Furthermore, defects in imprinted genes could cause severe diseases and abnormalities. Because of the importance of the ICRs to the regulation of parent-of-origin specific gene expression, I developed a genome-wide strategy for their localization. This strategy located clusters of the ZFBS-Morph overlaps along the entire human genome. Previously, I showed that in the mouse genome, clusters of 2 or more of these overlaps correctly located ∼ 90% of the fully characterized ICRs and germline Differentially Methylated Regions (gDMRs). The ZFBS-Morph overlaps are composite-DNA-elements comprised of the ZFP57 binding site (ZFBS) overlapping a subset of the MLL1 morphemes. My strategy consists of creating plots to display the density of ZFBS-Morph overlaps along genomic DNA. Peaks in these plots pinpointed several of the known ICRs/gDMRs within relatively long genomic DNA sections and even along entire chromosomal DNA. Therefore, peaks in the density-plots are likely to reflect the positions of known or candidate ICRs. I also found that by locating the genes in the vicinity of candidate ICRs, I could discover potential and novel human imprinting genes. Additionally, my exploratory assessments revealed a connection between several of the potential imprinted genes and human developmental anomalies including syndromes.

## INTRODUCTION

Imprinted genes play key roles in fetal development and in postnatal processes including behavior, sleep, feeding, maintenance of body temperature, metabolic regulation, and stem cell maintenance and renewal [2]. The imprinting mechanism is relatively complex and involves orchestrated actions of several enzymes and proteins; reviewed in [3]. Key players in that process include ZFP57 and a complex consisting of DNMT3A and DNMT3L. This protein-complex methylates DNA processively on a variety of CpG-rich substrates including the promoters of human genes encompassed by CpG islands [4]. Because of the importance of the ICRs to parent-of-origin specific gene expression, it is necessary to develop strategies for their localization in mammalian genomic DNA. Towards that goal, a previous study discovered that ZFP57 interacted with a CpG-methylated hexanucleotide (TGCCGC). This interaction was essential to the recognition of the ICRs by ZFP57 to maintain allele-specific gene expression [5]. Since CpG dinucleotides are infrequent in animal DNA [6], the instances of finding of ZFBS in mammalian DNA are by far less than those observed for AT-rich hexanucleotides. Nonetheless, since TGCCGC is relatively short it occurs often in genomic DNA [7]. Therefore, random occurrences of the hexameric site would obscure detection of the functional ZFP57 sites within the ICRs/gDMRs dispersed along relatively long genomic DNA sections.

Previously, I addressed that problem by extending the length of the canonical ZFP57 binding site to include additional nucleotides [7]. This strategy eliminated a significant fraction of randomly occurring ZFP57 sites in mouse genomic DNA and led to the discovery of the ZFBS-Morph overlaps [7, 8]. These overlaps define composite-DNA-elements consisting of the hexameric ZFP57 binding site overlapping a subset of the MLL1 morphemes [7, 8]. These morphemes correspond to the smallest ‘words’ in DNA that selectively bind the MT-domain in MLL1 [9]. The MT domain is present in MLL1 and MLL2. In DNA binding assays, this domain interacted selectively with nonmethylated CpG-rich sequences [10, 11]. Thus, within the ICRs, the ZFBS-Morph overlaps may play a dual and antagonistic role. Binding of MLL1 or MLL2 to ICRs would protect the DNA from methylation. Binding of ZPF57 to the modified DNA would maintain allele-specific expression [5, 7, 10-12].

Since closely-spaced ZFBS-Morph overlaps impart contextual specificity to ICRs, their localization could help with pinpointing the genomic positions of the ICRs that are currently unknown. Towards that goal, I describe a Bioinformatics strategy. This strategy involved creating plots to view the positions of clusters of 2 or more ZFBS-Morph overlaps along chromosomal DNA sequences. In these plots, peaks appear within a sliding window consisting of 850-bases. By displaying the plots at the UCSC genome browser, I could locate the peaks corresponding to several of the well-known ICRs/gDMRs and the peaks defining the genomic positions of candidate ICRs. I give an overview of how with the density-plots, I could discover potential imprinted genes. I found that several of these genes were associated with disease-states and developmental anomalies including syndromes. Examples included association of *IMPDH1* with Leber congenital amaurosis 11, *ARID1B* with Coffin-Siris syndrome, *PRDM8* with progressive myoclonic epilepsy-10, *PCNT* with microcephalic osteodysplastic primordial dwarfism type II, *CITED2* with ventricular septal defect 2, and *VAX1* with microphthalmia, cleft lip and palate, and agenesis of the corpus callosum.

## METHODS

### Marking the genomic positions of ZFP57 binding site and the ZFBS-Morph overlaps

From the UCSC genome browser, I downloaded the nucleotide sequences reported for the build hg19 of human chromosomes. Next, I created two texts files: one file containing the nucleotide sequences of the ZFBS-Morph overlaps [8]; the other the hexameric ZFP57 binding site [5]. Using a Perl script, initially I determined the genomic positions the ZFBS-Morph overlaps along the chromosomal DNA sequences. That script simultaneously opened the file containing the nucleotide sequence of a specified chromosome and the file containing the sequences of the ZFBS-Morph overlaps. After that step, the script moved along the DNA to report the genomic positions of the overlaps [8]. Next, I wrote a subroutine to combine the outputs obtained for various chromosomes. Another subroutine produced a file to create a custom track for displaying the genomic positions of the ZFBS-Morph overlaps at the UCSC genome browser. I followed similar procedures to obtain a file to display the genomic positions of the hexameric ZFP57 binding site at the UCSC genome browser [8]. Reference [13] gives a link for accessing the datafile containing the positions of the ZFBS-Morph overlaps and the hexameric ZFP57 binding site in the build hg19.

### Creating plots of the density of ZFBS-Morph overlaps in genomic DNA

I used another Perl script to obtain the genomic positions of DNA segments that covered 2 or more closely spaced ZFBS-Morph overlaps. That script opened the file containing the positions of ZFBS-Morph overlaps in a specified chromosome. Subsequently, the script scanned the file to count and to report the number of ZFBS-Morph overlaps within an 850-base window. By ignoring their isolated occurrences, the script removed background noise. Next, I combined and tailored the outputs of the program for display as a custom track at the UCSC genome browser. I chose the window-size by trial and error. Windows covering less than 850 bases tended to give superfluous spikes. Larger widows tended to produce false-peaks. In exploratory studies, I found that the density peaks corresponding to 3 or more ZFBS-Morph overlaps appeared reliable. Peaks covering 2 overlaps could be true or false-positive [14]. Reference [15] gives a link for accessing the datafile of the density-plots.

## RESULTS

### Robust peaks in the density-plots pinpointed several of the known ICRs/gDMRs

To investigate the validity of my strategy, initially I inspected the plot obtained for Chr6 to investigate whether I could locate the position of the ICR in the *PLAGL1* locus [16]. This ICR is ∼ 1 kb and maps to an intragenic CpG Island that encompasses the TSSs of a noncoding RNA gene (*HYMAI*) and several of the *PLAGL1* transcripts collectively known as *ZAC1* [16, 17]. In closeup views of the density-plots, apparent is a single robust intragenic peak mapping to the reported DNA segment that regulates parent-of-origin specific expression of *ZAC1* and *HYMAI* transcripts (Fig. 1).

**Figure 1.**
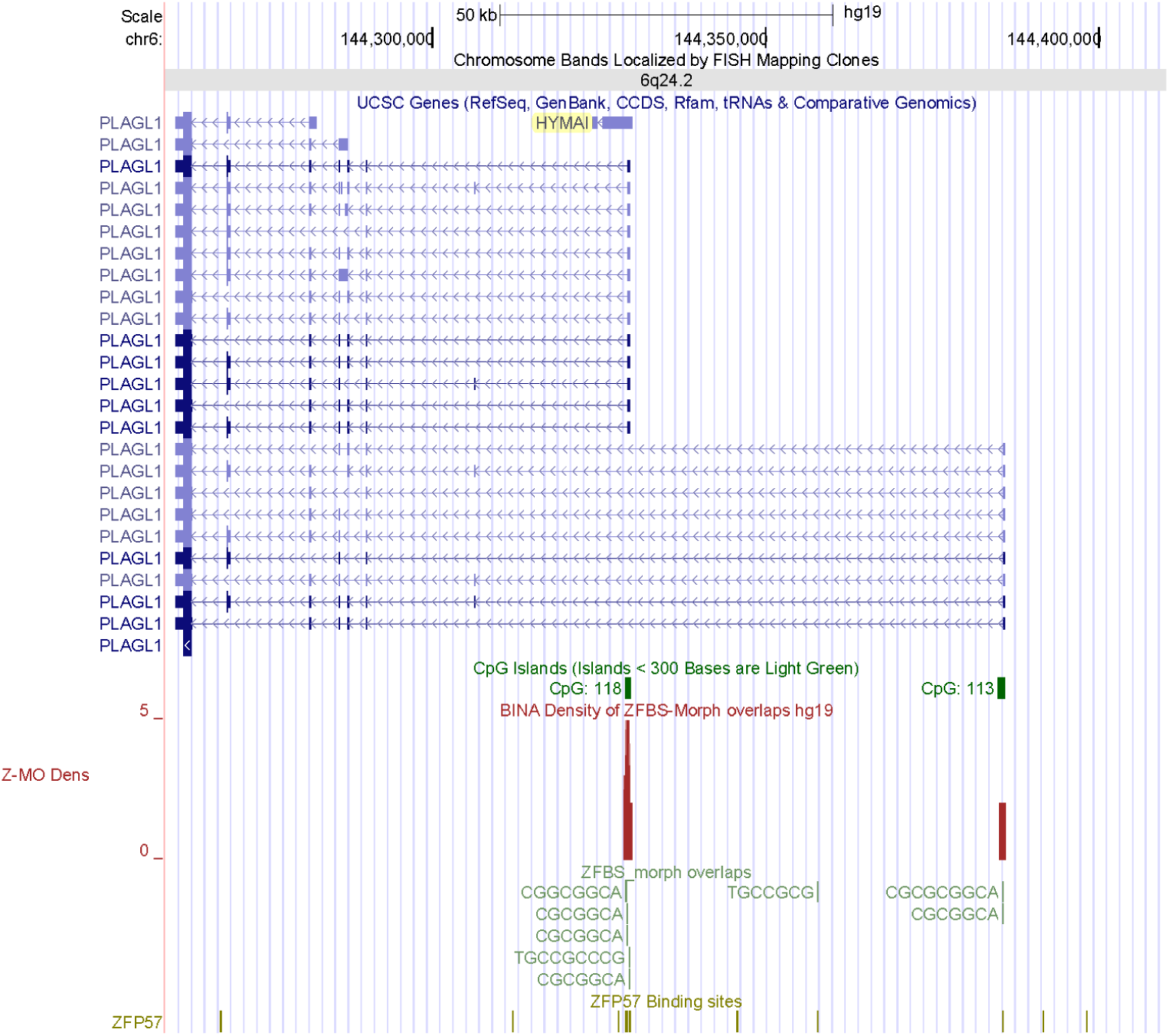
The position of a robust density peak locating the ICR in the *PLAGL1* locus. In descending order, tracks display the positions of the chromosomal bands (gray), genes and transcripts (blue), CGIs (green), peaks in the density-plots (maroon), the sequences of ZFBS-Morph overlaps shown in pack format (hunter green), the canonical ZFP57 in dense format (olive green).

Next, I inspected the density-plot of Chr11 to investigate the positions of peaks with respect to two well-known imprinted domains. The telomeric/distal domain 1 includes a noncoding RNA gene (*H19*) and two genes encoding insulin-like growth factor 2 (*IGF2*) and insulin (*INS*). Domain 2 is relatively long and encompasses three noncoding RNA genes (*KCNQ1OT1, KCNQ1-AS1* and, *KCNQ1DN*) and ∼ ten protein coding genes including *KCNQ1, CDKN1C, SLC22A18, PHLDA2*, and *OSBPL5* [18]. My initial assessments of plots included several long DNA sections encompassing both the *H19* – *IGF2* and the *KCNQ1* imprinted domains. In a 1.4 Mb long DNA, I observed 3 robust density peaks and several peaks covering 2 ZFBS-morph overlaps (Fig. 2). One of the robust peaks appears as a doublet and maps to the imprinted domain 1. The other two map to imprinted domain 2. In imprinted domain 1, a single gDMR/ICR regulates transcription of *H19* selectively from the maternal allele and expression of *IGF2* and *INS* from the paternal allele [19]. The ICR in domain 1 is upstream of *H19* TSS and is often described in the context of several unique repeats [20] and sites predicted to bind the transcription factor CTCF [21]. Among these sites, the ENCODE data do not support the existence of CTCF site 5; for details see reference [22]. In closeup views, I observed that the DNA section encompassing the doublet, correctly located the ICR in the imprinted domain 1 (Fig. S1 shown after the references).

**Figure 2.**
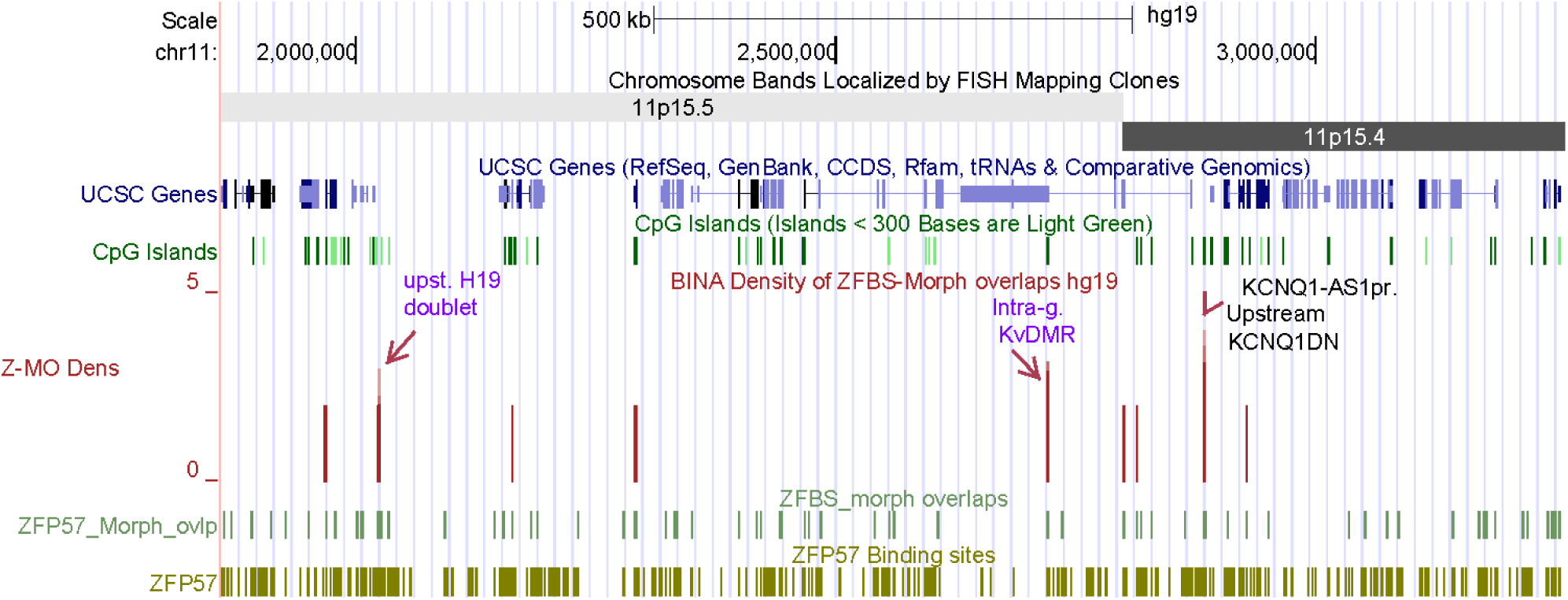
The positions of peaks in a density-plot covering 1.4 Mb long DNA. The density-plot is shown in full format. The other tracks are displayed in dense-format. Purple letters denote the known ICRs. Black letters denote an intergenic candidate ICR.

Telomeric with respect to domain 1, domain 2 is relatively long and includes the ICR (KvDMR1) that regulates expression of several imprinted genes [18, 23]. This intragenic ICR is in the vicinity of *KCNQ1OT1* TSS. In closeup views, I observed a robust density peak within a CpG island encompassing the TSS of *KCNQ1OT1.* Thus, this peak correctly located the KvDMR1 (Fig. S2). Closeup views also revealed another robust density peak between the two noncoding RNA genes corresponding to *KCNQ1DN* and *KCNQ1-AS1*. One of the noncoding RNA genes (*KCNQ1DN*) corresponds to an imprinted transcript. It is located within the WT2 critical region [24]. The other gene (*KCNQ1-AS1*) was observed in the reference genome assembly (https://www.ncbi.nlm.nih.gov/nuccore/NR_130721.1). Even though the function of *KCNQ1-AS1* is unknown, it is worth noting that it is transcribed antisense with respect to *KCNQ1OT1*. Therefore, the expression of *KCNQ1-AS1* might be a mechanism to impede leaky production of *KCNQ1OT1* in a subset of human tissues.

### The density -plots predicted ICRs for parent-of-origin specific expression of several experimentally identified candidate imprinted genes

To further explore the robustness of my approach, I inspected the positions of the density peaks with respect to the genes listed in Table 4 of a report that analyzed tissues from human term placenta using high-throughput analyses [25]. That list included 39 candidate imprinted genes. For evaluations, I analyzed the density-plots to investigate whether these genes also corresponded to potential imprinted genes predicted by my approach. Among the entries in that list, the UCSC genome browser could not find *MGC16597* and *MGC24665*. Among the remaining candidates, I found density peaks within or in the vicinity of *PRDM8, SQSTM1, NM_006031, TJP2, CDK2AP1, MYH7B*, and *MAN2C1*.

In the *PRDM8* locus, I observed a robust intragenic density peak. This peak maps to a CGI in the last exon of several *PRDM8* transcripts (Fig. S3). This finding agrees with the correspondence of *PRDM8* to an actual imprinted gene. Next, I examined the positions of density peaks in a DNA segment that included *SQSTM1*. This segment contains two overlapping genes (*MGAT4B* and *SQSTM1*). A relatively long CpG island encompasses the TSSs of three *SQSTM1* transcripts and the longest *MGAT4B* transcript (Fig. 3). I observed an intragenic density peak within that island. Thus, the density-plots revealed a candidate ICR regulating parent-of-origin specific expression of *SQSTM1*. Furthermore, while lending support for the correspondence of *SQSTM1* to a genuine imprinted gene, the density-plots located a potential imprinted transcript produced from *MGAT4B* (Fig. 3). For the gene listed as *NM_006031* in reference [25], the UCSC genome browser displayed the corresponding gene (*PCNT*). Within that locus, the density-plots revealed two very robust peaks supporting that *PCNT* could be a genuine imprinted gene (Fig. S4). For four of the listed loci (*TJP2, CDK2AP1, MYH7B*, and *MAN2C1*), primarily I observed peaks covering 2 ZFBS-Morph overlaps (plots not shown). These peaks could be true or false positives [14]. For *FLJ10300*, the genome browser showed a sequence (AK001162) mapping to *WDR60*. Within that gene, I observed a density peak covering 2 ZFBS-Morph overlaps (Fig. S5). However, in plots I noticed a very robust density peak mapping to the 5’ end of *ESYT2* (Fig. S5). This peak could a better candidate ICR for regulating parent-of-origin expression of *WDR60*. For several of the listed candidate imprinted genes [25], I noticed density peaks far upstream or downstream of their TSSs. Examples include *XRRA1, CD151*, and *VPS11* (plots not shown). Thus, it seemed that density peaks could locate potential ICRs for several candidate imprinted genes discovered by experimental and computational strategies.

**Figure 3.**
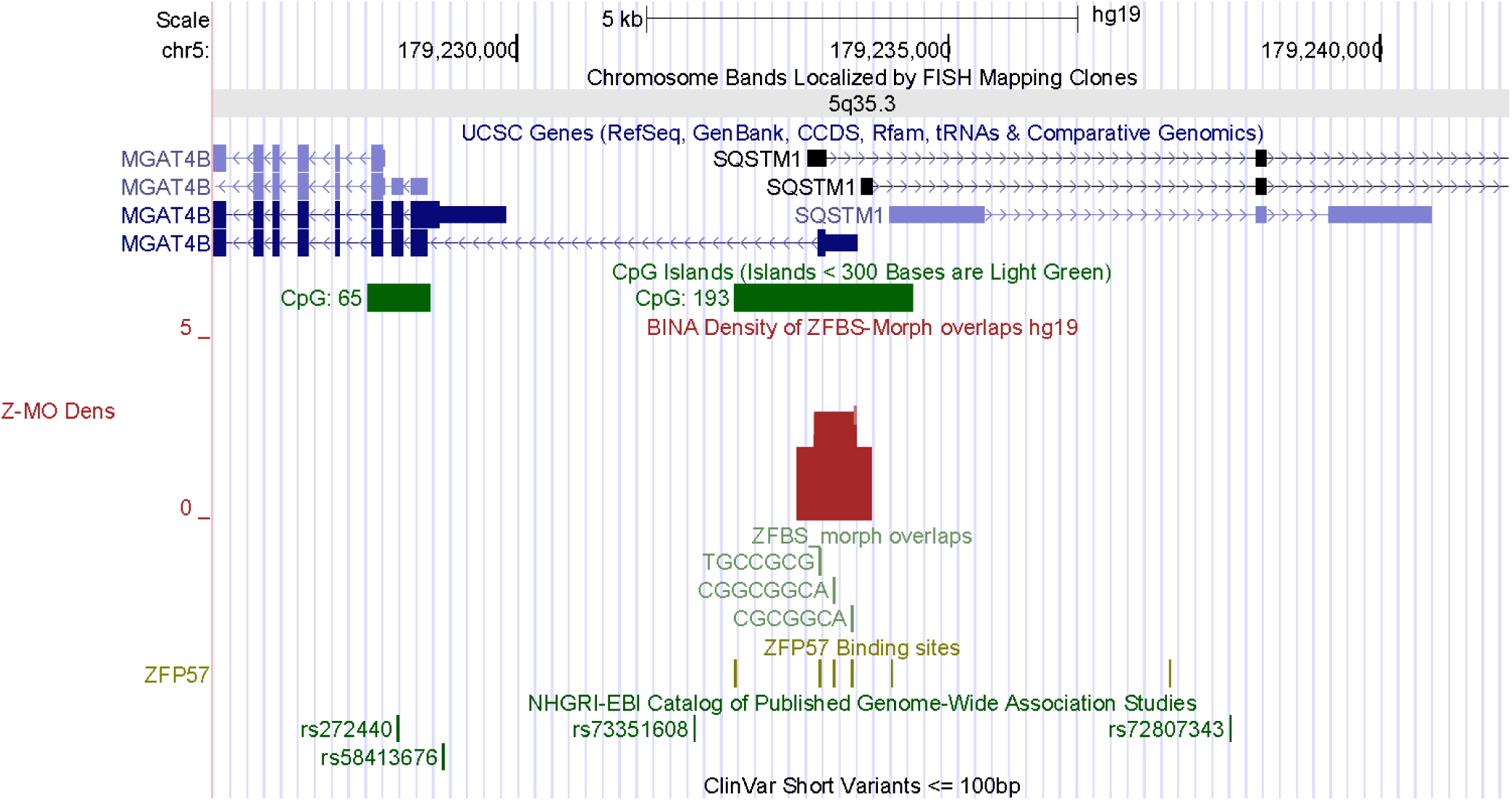
A candidate ICR mapping to overlapping transcripts (*MGAT4B* and *SQSTM1*). High-throughput experimental analysis identified *SQSTM1* as a candidate imprinted gene [25]. The density-plots include a peak corresponding to a candidate ICR for imprinted expression of *SQSTM1*. Furthermore, the position of this ICR predicts parent-of-origin specific expression for the longest transcript produced from *MGAT4B*.

### The density-plots offer a strategy to discover candidate ICRs and potential imprinted genes, and to determine whether these genes could be associated with clinical abnormalities

In humans, defects in imprinted genes could cause severe diseases and genetic anomalies [2]. Therefore, in my studies, I explored whether the density-plots could help with finding the unknown imprinting genes and whether any of these genes could be associated with known genetic disorders. In initial exploratory studies, I examined the positions of peaks with respect to short variants along the entire Chr6 (Fig. 4). In maps, I noticed that these variants were dispersed along the entire chromosome. To examine whether I could find a connection between the potential imprinted genes and genetic abnormalities, I sampled and inspected several large DNA sections that contained robust density peaks.

**Figure 4.**
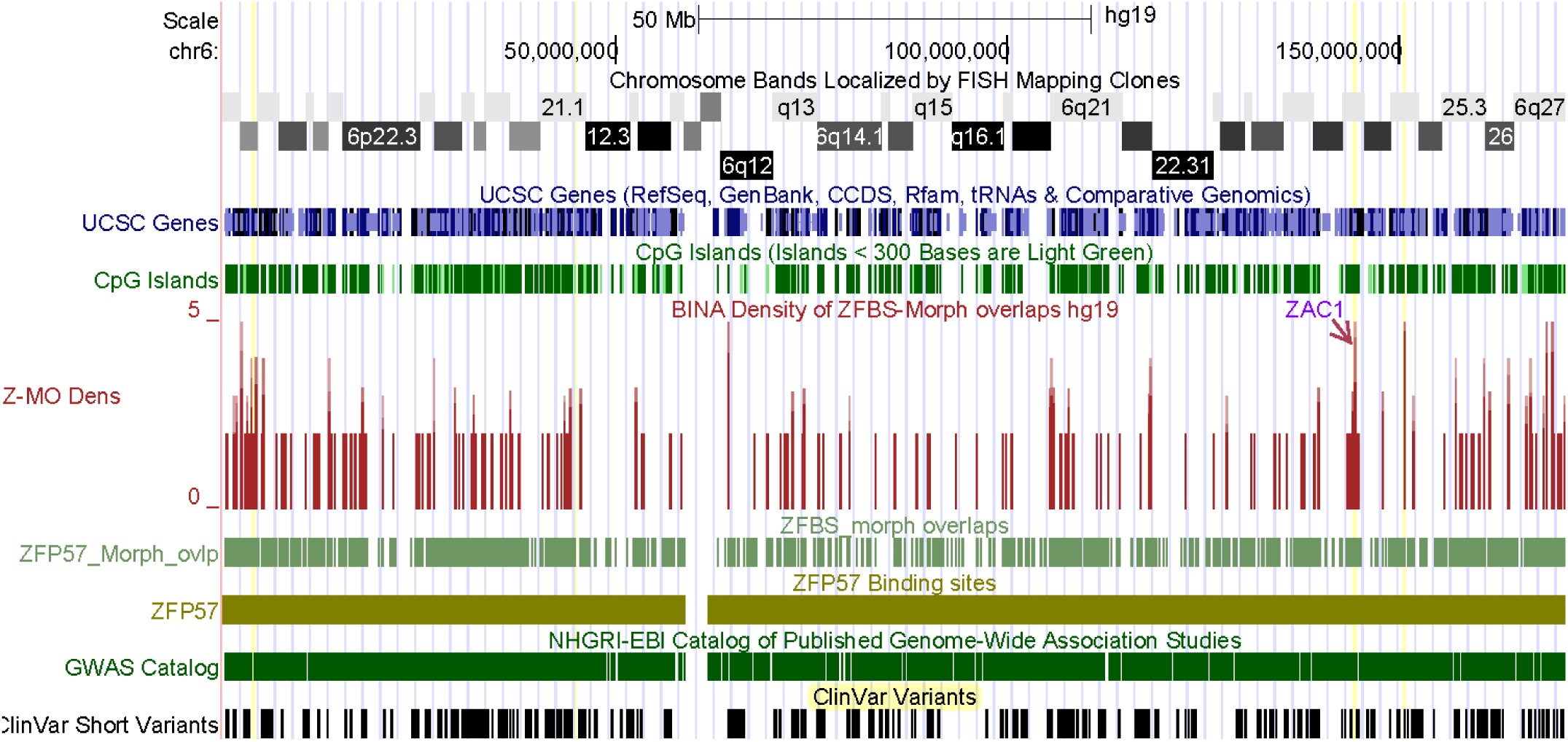
Density-plot displaying the positions of peaks with respect to GWAS studies and clinical variants along Chr6. With such plots, it is possible to view the positions of a significant fraction of the known and candidate ICRs along an entire chromosome. Furthermore, my approach facilitates examining the potential imprinted genes in the context of diseases and genetic abnormalities.

### A DNA section from Chr6q contains a known ICR and several candidate ICRs for potential imprinted genes

For inspections, initially I selected a 25 Mb long DNA encompassing several chromosomal bands (Fig. 5). Even within such a long genomic DNA section, the density peaks appear clearly resolved. In the displayed view, I observed 7 robust density peaks (1 per ∼ 3.6 Mb) demonstrating that the robust peaks occurred sporadically in human genomic DNA. One of the robust peaks pinpointed the known intragenic ICR of *ZAC1* and *HYMAI* (Fig. 5). The remaining 6 reflect the positions of candidate ICRs dispersed in various genomic locations.

**Figure 5.**
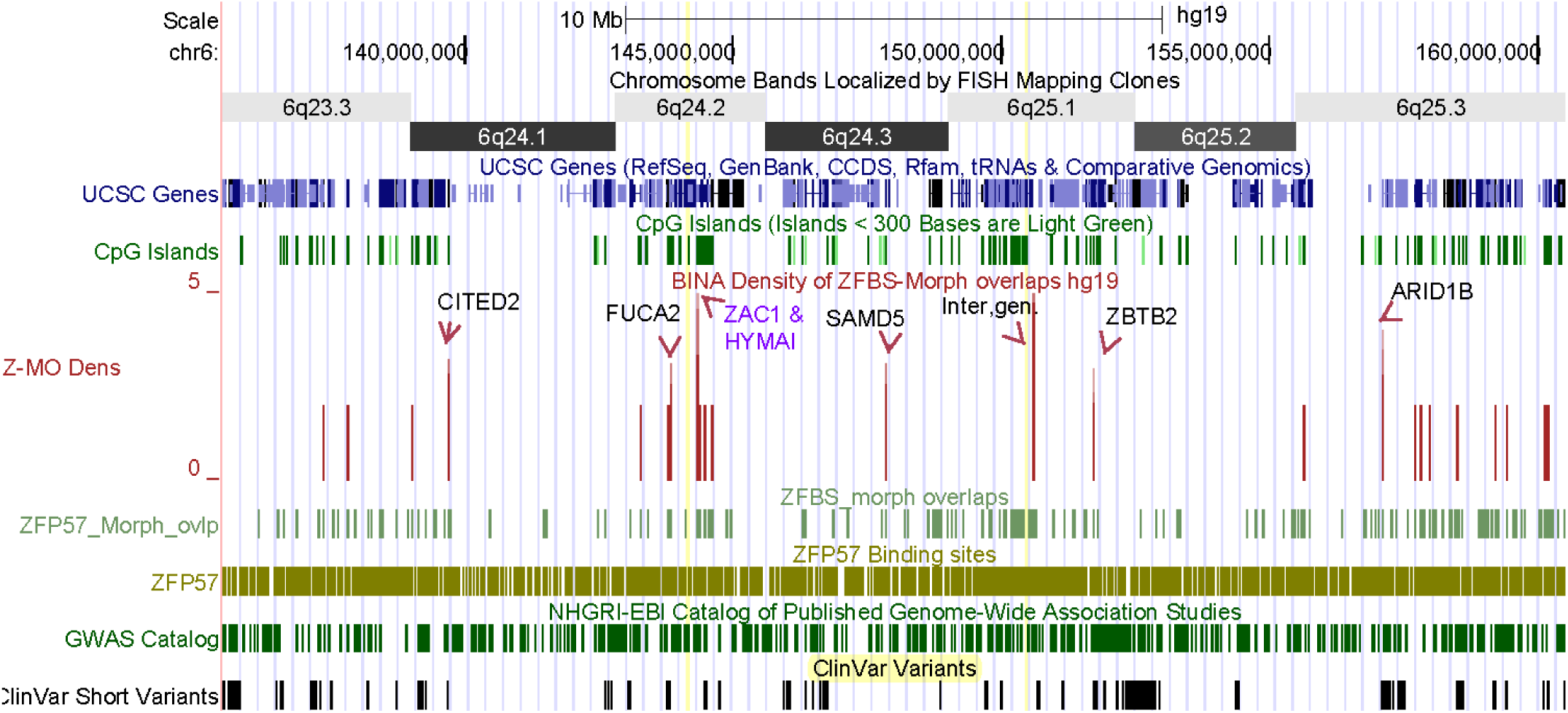
Discovering candidate ICRs and novel imprinting genes. The displayed DNA section includes several density-peaks dispersed along chromosomal bands. One of the peaks maps to the ICR of *ZAC1* and *HYMAI*. The remaining robust peaks correspond to candidate ICRs for potential imprinted genes or transcripts.

Chr6q25.1 includes an intergenic candidate ICR between *PPP1R14C* and *IYD*. The other candidates are dispersed along the 25 Mb DNA mapping to *CITED2, FUCA2, SAMD5, ZBTB2*, and *ARID1B* (Fig. 5). The candidate ICR for *CITED2* is in Chr6q24.1. Its corresponding density peak maps to an intragenic CpG island in the last *CITED2* exon (Fig. S6). CITED2 (CBP/p300-interacting transactivator 2) regulates transcription. Absence of *Cited2* in mouse embryos caused congenital heart disease by perturbing left-right patterning of the body axis [26]. In Chr6q24.2, a candidate ICR maps to an intragenic CpG island near the 5’ end of the longest of *FUCA2* transcript (Fig. S7). Genome-wide analyses have identified *FUCA2* and *IL-18* as novel genes associated with diastolic function in African Americans with sickle cell disease [27]. In chr6q24.3, a candidate ICR maps to a CpG island that encompasses *SAMD5* TSS (Fig. S8). A study found that *SAMD5* was overexpressed in prostate cancer and had powerful prognostic ability for predicting post-operative biochemical recurrence after radical prostatectomy [28].

In Chr6q25.1, a candidate ICR maps to a CpG island at the 5’ end of *ZBTB2* (Fig. S9). ZBTB2 binds DNA and is among the master regulators of the p53 pathway [29]. In mouse embryonic stem cells, ZBTB2 dynamically interacted with nonmethylated CpG island promoters and regulated differentiation [30]. In colorectal cancer, the abnormal forms of *ZBTB2* increased cell proliferation [31]. Chr6q25.3 includes 1 candidate ICR (Fig. 5). This ICR corresponds to 2 density peaks (Fig. 6). One peak encompasses *ARID1B* TSS. The other maps to the *ARID1B* 1^st^ exon. *Arid1b* is a known imprinted gene in mouse [1]. Therefore, from my data, one could deduce that its human ortholog also is an imprinted gene. *ARID1B* encodes an enzyme that removes H3K4 methyl-marks from chromatin [32]. In mouse ES cells, depletion of *Arid1b* caused defects in gene expression programs [33]. In Chr6q25.1, I observed a candidate ICR between *PPP1R14C* and *IYD* (Fig. 7). This peak is in a DNA segment in the vicinity of the 3’ end *PPP1R14C* and far upstream of *IYD* TSS. PPP1R14C regulates the enzymatic activity of protein phosphatase 1. *IYD* encodes an enzyme that functions in iodide salvage in the thyroid [34]. Notably, the thyroid hormone pathway includes another enzyme (DIO3) that is encoded by an imprinted gene [35].

**Figure 6.**
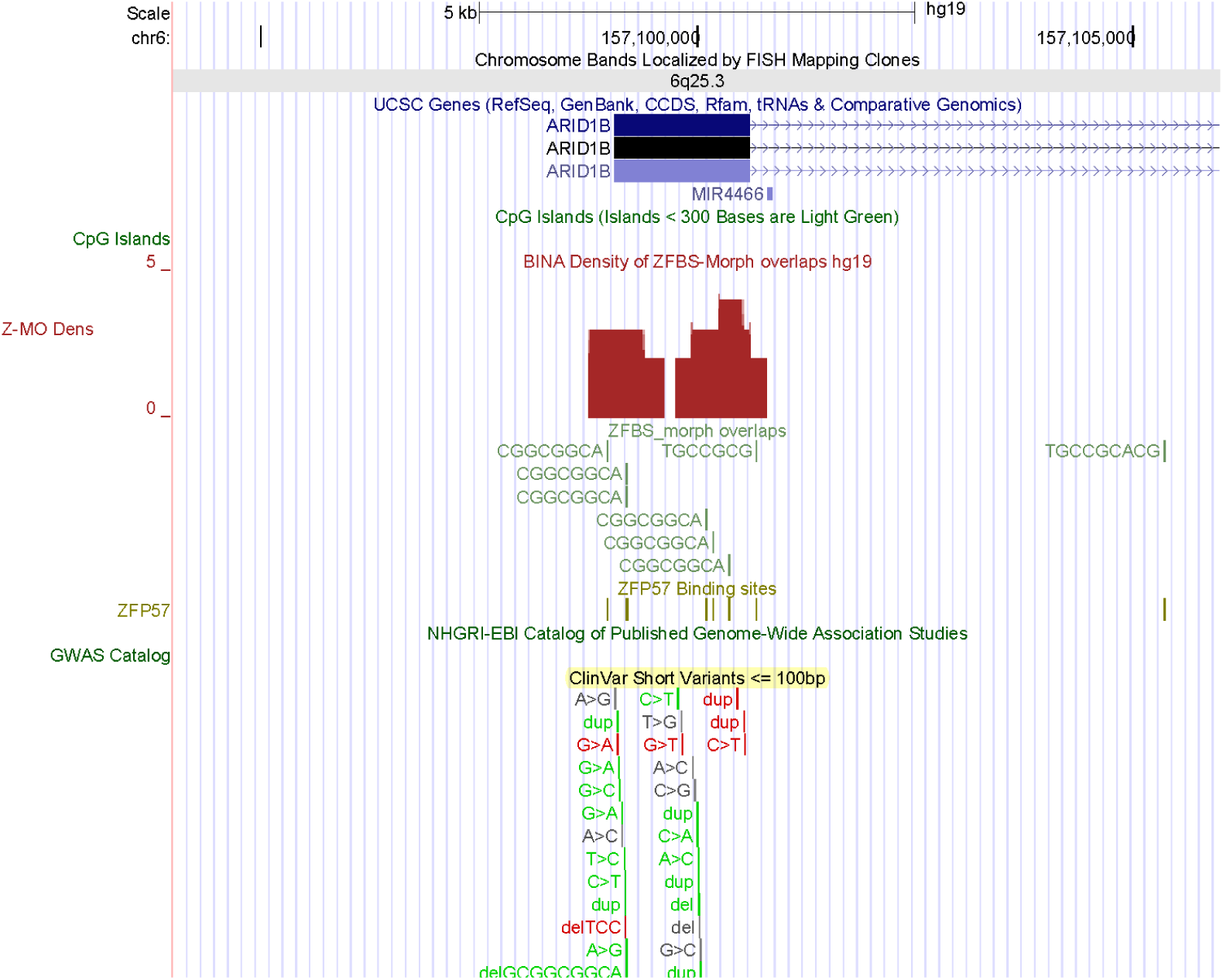
A candidate ICR regulating parent-of-origin specific expression of *ARID1B*. In mouse, *Arid1b* is a known imprinted gene [1].

**Figure 7.**
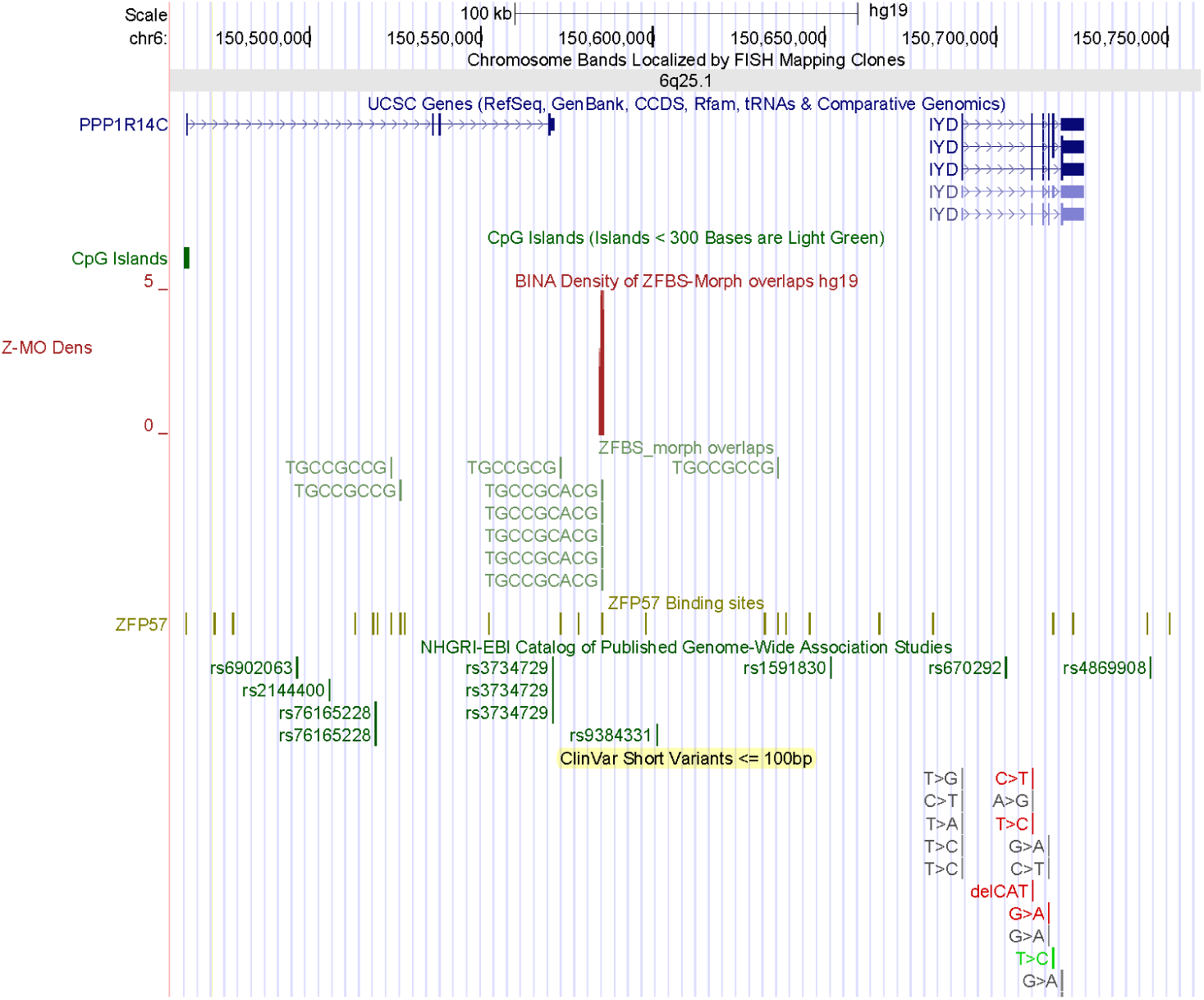
An intergenic candidate ICR regulating parent-of-origin specific expression of two potential imprinted genes (*PPP1R14C* and *IYD)*.

### A DNA section from Chr7q includes the known ICR of MEST, a candidate ICR for regulating expression from a known imprinted gene (KLF14), and a candidate ICR for a potential imprinted gene (IMPDH1)

Chr7 contains several known imprinted genes [36, 37]. From Chr7qA, I selected a 4 Mb long DNA covering 3 robust density peaks (1 robust peak per 1.3 Mb). These peaks map to *IMPDH1, MEST*, and *KLF14* (Fig. 8). The *MEST* and *KLF14* loci includes known imprinted transcripts [38]. In the *MEST* locus, an intragenic ICR regulates parent-of-origin-specific expression of a subset of *MEST* transcripts and *MESTIT1*, a noncoding RNA gene [39, 40]. An enlarged view of the density-plots shows a peak within an intragenic CpG island that encompasses the TSSs of both *MEST* and *MESTIT1* (Fig. S10). Thus, this peak correctly located the ICR in the *MEST* locus.

**Figure 8.**
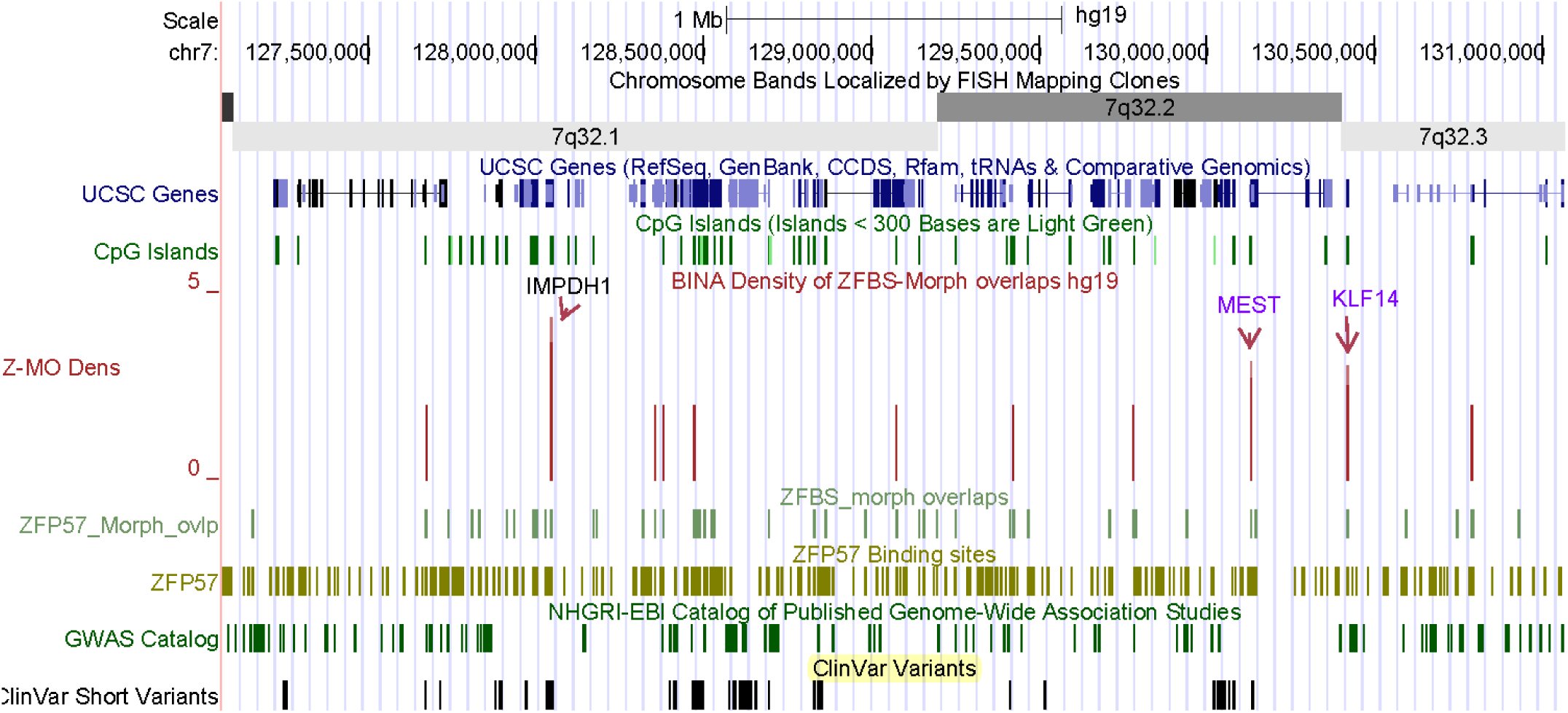
A long DNA section containing the ICR in the *MEST* locus, a known imprinting gene (*KLF14*), and a candidate ICR for a potential imprinted gene (*IMPDH1*).

The displayed view also shows the peak that corresponds to *KLF14* (Fig. S10). This gene is imprinted in human but not in mouse. *KLF14* is the first example of an imprinted gene that has undergone accelerated evolution in the human lineage [41]. To date, I have not found a report locating the ICR of *KLF14*. Notably, density plots revealed a candidate ICR within a CpG island encompassing *KLF14* TSS (Fig. 8). Hence, this candidate ICR might regulate parent-of-origin specific expression of *KLF14* in human DNA.

Next, I obtained a closeup view to inspect the position of the candidate ICR that mapped to *IMPDH1* (Fig. 9). As observed for the *MEST* locus, this ICR is intragenic and maps to a CpG island that encompasses the TSSs of several of short *IMPDH1* transcripts (Fig. 9). Even though *IMPDH1* is expressed in many tissues, its predominant transcripts are produced in the inner segment and synaptic terminals of retinal photoreceptors [42]. The IMPDH proteins form active homo-tetramers that catalyze the rate-limiting step for *de novo* guanine synthesis by converting inosine monophosphate to xanthosine monophosphate [42]. Deleterious mutations in *IMPDH1* is associated with Leber congenital amaurosis 11 early-onset childhood retinal dystrophies (https://www.omim.org/entry/613837).

**Figure 9.**
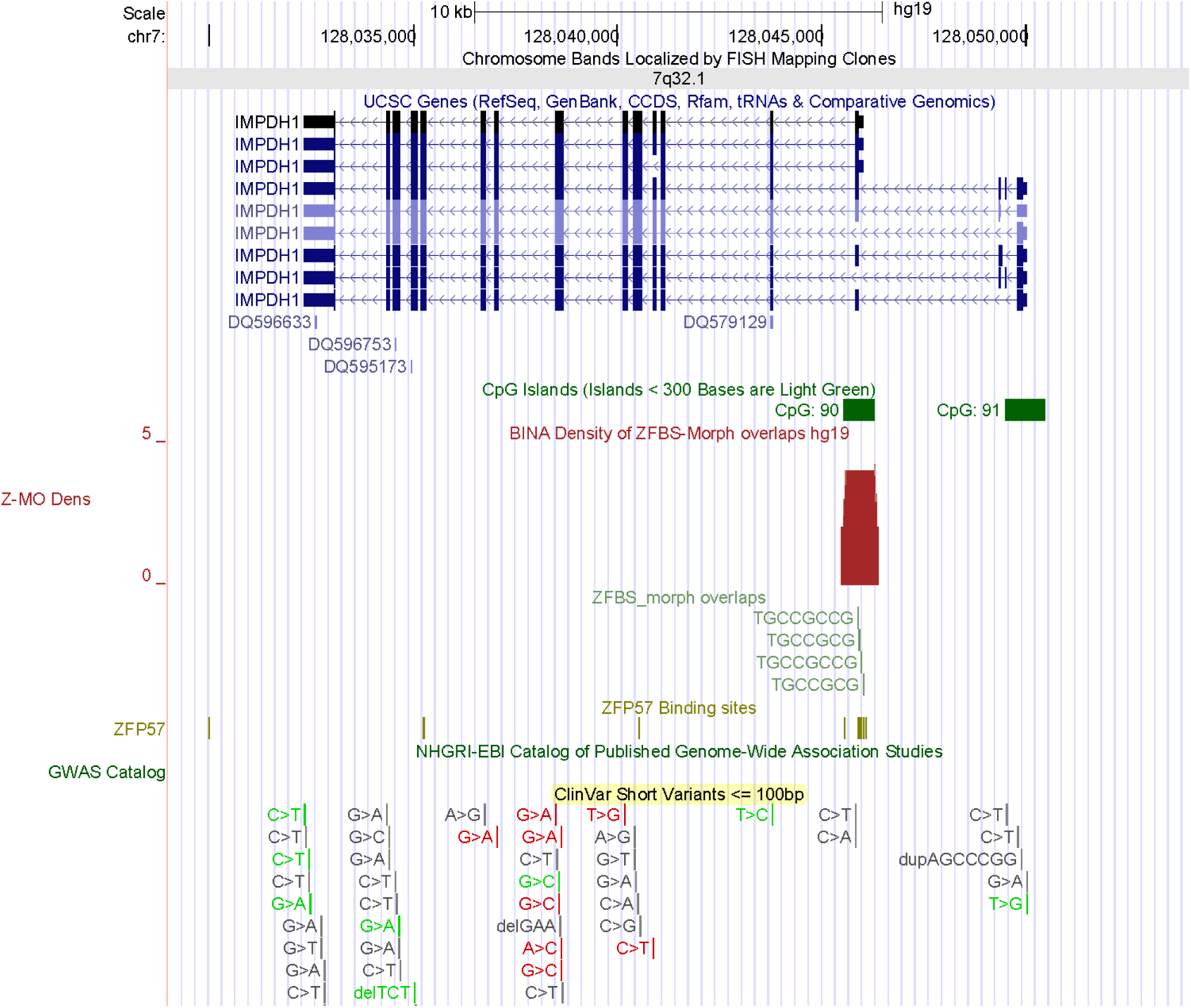
A candidate ICR mapping to a CGI that encompasses the TSS of several intragenic *IMPDH1* transcripts.

### A long DNA section from Chr10q includes the known ICR of INPP5F_v2 and a candidate ICR for VAX1, a potential imprinted gene

In Chr10q, a nearly 8.6 Mb DNA encompasses several chromosomal bands and 2 robust density peaks (1 per 4.3 Mb). One of the peaks maps to the *INPP5F* locus in Chr10q26.11. The other maps to *VAX1* in 10q25.3 (Fig. 10). From *INPP5F* are produced several transcriptional variants. In mouse, one of the variants (*Inpp5f_v2*) is imprinted in the brain [43]. The TSS of *Inpp5f_v2* is within a differentially methylated CpG island [43]. Allele-specific expression was observed for both mouse and orthologous human *INPP5_v2* transcript [43, 44]. In closeup views of the density-plots, I observed an intragenic density peak that pinpointed the ICR for parent-of-origin specific expression of human *INPP5_v2* (Fig. S11). The density peak in the other locus provides a candidate ICR for a potential imprinted gene (*VAX1*). This gene encodes a transcription factor with a homeobox for binding DNA. The candidate ICR maps to a CpG island that encompasses TSSs of 2 *VAX1* transcripts (Fig. 11). In mouse, *Vax1* is expressed in the pituitary, hypothalamus, and testis [45]. A study implicated two homozygous mutations in *VAX1* causing microphthalmia associated with cleft lip and palate and agenesis of the corpus callosum [46].

**Figure 10.**
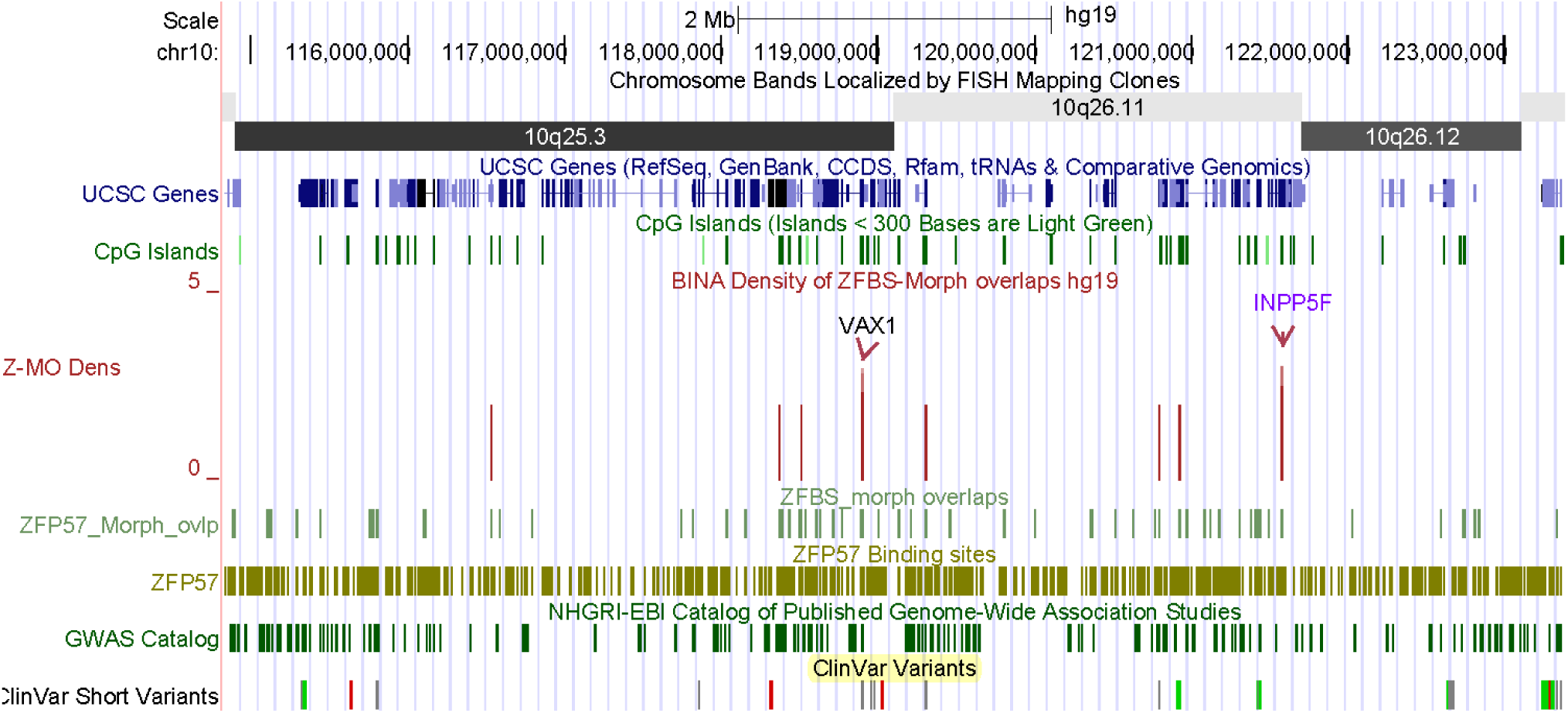
A long DNA section includes the known ICR in the *INPP5F* locus and a candidate ICR for a potential imprinted gene (*VAX1*).

**Figure 11.**
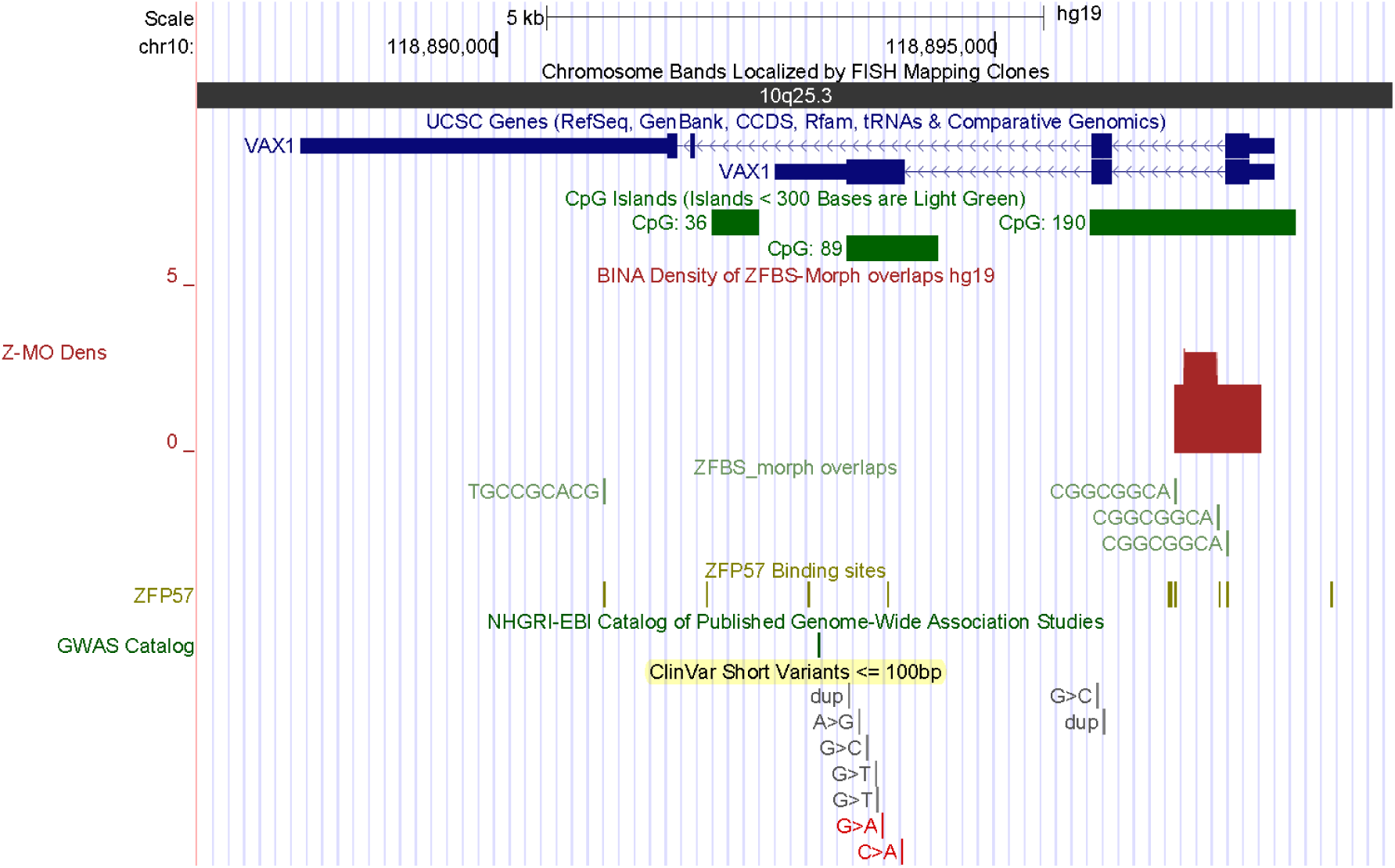
The candidate ICR regulation parent-of-origin specific expression of a potential imprinted gene (*VAX1*).

## DISCUSSION

The discovery of unknown ICRs would facilitate pinpointing the genes in their vicinity and thus uncovering novel imprinted genes. Furthermore, examination of the imprinted genes in the context of clinical variants gives clues into their impacts on human embryonic development, disease states, and genetic anomalies including syndromes [2]. However, to date I could not find any genome-wide studies to methodically discern the genomic positions of the ICRs and to obtain a nearly complete list of novel imprinted genes and transcripts. Towards these and related goals, I developed a predictive genome-wide strategy. My strategy pinpointed several of the known ICRs within relatively long DNA sections. Examples include the ICRs of *H19* – *IGF2* and *KCNQ1* imprinted domains in 1.4 Mb long DNA (Fig. 2); the ICR of *MEST* and *MESTIT1* transcripts in 4.0 Mb DNA (Fig. 8), and the ICR of *NPP5F_v2* within a nearly 8.6 Mb DNA (Fig. 10). Even along the entire Chr6, I could discern the ICR of *ZAC1* and *HYMAI* in the *PLAGL1* locus (Fig. 4).

My strategy involves creating density-plots to detect the genomic DNA segments that contain clusters of 2 or more ZFBS-morph overlaps [13, 15]. Previously, I showed that occurrences of such clusters pinpointed ∼ 90% of the fully characterized ICRs/gDMRs in the mouse genome [7, 8, 47]. Even though my approach is predictive, it is based on reports demonstrating the importance of ZFP57 in maintaining allele-specific gene repression [5, 12, 48, 49]. Furthermore, my discovery of ZFBS-morph overlaps has offered mechanistic clues into why ZFP57 is selectively recruited to the ICRs but not elsewhere in the genomic DNA. Briefly, essential to genomic imprinting is a protein complex consisting of DNMT3A and DNMT3L [50]. This complex methylates DNA processively [4]. Since clusters of ZFBS-morph overlaps are CpG-rich, they could provide sites for DNMT3A-DNMT3L to methylate several CpGs processively and thus facilitating the propagation step of DNA methylation by other DNMTs. Furthermore, the methylated ZFBS-morph overlaps would recruit ZFP57 to associate selectively with the ICRs to maintain parent-of-origin specific gene repression [7].

### The majority of candidate ICRs maps to CpG islands; a subset maps to specific gene transcripts

The UCSC genome browser is highly suitable for examining results of predictive methods in the context of landmarks including the positions of chromosomal bands, genes, transcripts, and the CpG islands [51]. The islands that include ICRs could be intergenic, encompass promoters, TSSs, and the 1^st^ exon of genes [52]. The ICRs also occur in intragenic CpG islands [39, 53, 54]. Similarly, in the density-plots the analyzed robust peaks primarily map to CpG islands at various genomic locations. Examples include the candidate ICRs for *PPP1R14C, IYD, CITED2, IMPDH1*, and *VAX1* (Figs. 7, S6, 8, 9, and 11). Several of the known ICRs correspond to single transcripts or to transcriptional variants [52]. I observed similar patterns for several of the candidate ICRs. For example, the candidate ICRs for *SAMD5* and *ZBTB2* correspond to isolated transcripts (Figs. S8 and S9). The candidate ICR for *FUCA2* corresponds to the longest transcriptional variant (Fig. S7). The candidate ICR for *IMPDH1* encompasses intragenic TSSs (Fig. 9).

### Density-plots revealed candidate ICRs for potential imprinted genes associated with disease states and syndromes

With animal model systems, it is possible to determine whether knockout of a gene would produce a phenotype. In humans, one could examine adverse effects of anomalous loci in the context of the clinical variants that produce discernable phenotypes. Examples include developmental disorders, neurological disorders, malformation of body parts, and syndromes. One could identify these phenotypes from literature surveys, the track displaying the clinical variants at the UCSC browser, or both. For example, examine the figure that displays the positions of short clinical variants with respect to peaks in the density-plot obtained for the entire Chr6 (Fig. 4). As these peaks, the clinical variants are primarily dispersed in gene-rich genomic DNA sections.

*ARID1B* is among the potential human imprinted genes identified by my approach (Fig. 6). In mouse, *Arid1b* is a known imprinted gene [1]. My data predicts that human *ARID1B* also is an imprinted gene. This gene encodes an enzyme that removes activating H3K4 methyl marks from chromatin to repress transcription [55]. In human, deleterious variations in *ARID1B* are thought to contribute to Coffin-Siris syndrome. This abnormality is a multiple malformation syndrome characterized by mental retardation associated with coarse facial features, hypertrichosis, sparse scalp hair, and hypoplastic or absent fifth fingernails or toenails. Other features may include poor overall growth, craniofacial abnormalities, spinal anomalies, and congenital heart defects (https://www.omim.org/entry/135900). Mechanistically, within the protein-networks ARID1B interacts with a complex Known as NuRD (Nucleosome Remodeling and Deacetylase); for details see Fig. 1 in reference [7]. In the networks, NuRD is a central node for receiving or transmitting signals via protein-protein interactions. For example, while one of the subunits in NuRD (Mi-2α) interacts with TRIM28, its HDAC1 subunit interacts with MLL1, DNMT3A, DNMT3L, and H3K4 demethylases including ARID1B [7].

Several of the potential imprinted genes identified by my approach correspond to a subset of candidate-imprinted genes discovered by experimental techniques [25]. Examples include *SQSTM1, PRDM8*, and *NM_006031*/*PCNT* (Figs. 3, S3, and S4). A domain in PRDM8 methylates H3K9 and thus impacts the chromatin structure [56]. Monoallelic expression of *PRDM8* was detected in placental tissues [25]. Genetic studies have observed association of abnormality in *PRDM8* with progressive myoclonic epilepsy-10. This anomaly is an autosomal recessive neurodegenerative disorder characterized by onset of progressive myoclonus, ataxia, spasticity, dysarthria, and cognitive decline in the first decade of life (https://www.omim.org/entry/616640). PCNT (pericentrin) is a component of pericentriolar material that surrounds the two centrioles of a centrosome [57]. Absence of *PCNT* results in disorganized mitotic spindles and missregulation of chromosomes [58]. Microcephalic osteodysplastic primordial dwarfism type II (MOPD2) is caused by homozygous or compound heterozygous mutations in the PCNT gene (https://www.omim.org/entry/210720). MOPD2 is characterized by intrauterine growth retardation, severe proportionate short stature, and microcephaly. Adults with this inherited disorder have an average height of 100 centimeters and a brain size comparable to that of a 3-month-old baby. Otherwise, they have near-normal intelligence [58].

Potential imprinted genes discovered *de novo* by my approach include *IMPDH1* (Fig. 9), *CITED2* (Fig. S6), *SAMD5* (Fig. S8), *ZBTB2* (Fig. S9), and *VAX1* (Fig. 11). IMPDH1 (inosine monophosphate dehydrogenase 1) catalyzes the synthesis of xanthine monophosphate [59]. This reaction is the rate-limiting step in the *de novo* synthesis of guanine nucleotides. Several of the deleterious mutations in *IMPDH1* are associated with Leber congenital amaurosis 11 (LCA11). This anomaly consists of a group of early-onset childhood retinal dystrophies characterized by vision loss, nystagmus, and severe retinal dysfunction (https://www.omim.org/entry/613837). Mutations in *IMPDH1* also may cause retinitis pigmentosa-10 (RP10). In most patients, this anomaly is characterized by early onset and rapid progression of ocular symptoms, beginning with night blindness in childhood, followed by visual field constriction (https://www.omim.org/entry/180105).

CITED2 transactivates transcription through interactions with CBP/P300 [60]. Deleterious mutations in *CITED2* cause ventricular septal defect 2 (VSD2). This disorder is the most common form of congenital cardiovascular anomaly. It occurs in nearly 50% of all infants with a congenital heart defect and accounts for 14% to 16% of cardiac defects that require invasive treatment within the first year of life (https://www.omim.org/entry/614431). Congenital VSDs may occur alone or in combination with other cardiac malformations. In that context, it seems relevant that loss of *Cited2* in mouse causes congenital heart disease by perturbing left-right patterning of the body axis [26]. Therefore, cardiac malformations in VSDs might in part arise from defects in left-right patterning of the body axis.

In the context of disease-related anomalies, a study found that *SAMD5* was overexpressed in prostate cancer and had powerful prognostic ability on predicting post-operative biochemical recurrence after radical prostatectomy [28]. *ZBTB2* is among the genes found in approximately 15% of all colorectal cancers [31]. In colorectal cancer, The abnormal forms of *ZBTB2* increased cell proliferation [31]. This gene encodes a transcription factor. Several proteins in the ZBTB family have emerged as critical factors that regulate the lineage commitment, differentiation, and function of lymphoid cells as well as many other developmental events [61]. ZBTB2 is among the master regulators of the p53 pathway [29]. In mouse embryonic stem cells, ZBTB2 dynamically interacted with unmethylated CpG island promoters and regulated differentiation. *VAX1* also encodes a transcription factor that controls developmental processes. The structure of VAX1 includes a homeodomain for binding DNA. In mouse, this homeobox-containing gene is expressed in the developing anterior ventral forebrain [62]. In mouse, *Vax1* is expressed in the pituitary, hypothalamus, and testis [45]. From studies of 70 patients, a report found associations of two homozygous mutations in *VAX1* with microphthalmia, cleft lip and palate, and agenesis of the corpus callosum [46]. For an overview, see (https://www.omim.org/entry/614402).

## CONCLUSION

To researchers, I have offered a predictive genome-wide strategy to discover candidate ICRs and novel imprinted genes. I gave evidence for robustness of my strategy by pinpointing several of the well-known ICRs in relatively long DNA sections. I also gave examples showing that my strategy predicted ICRs for several of the candidate imprinted genes discovered by experimental strategies. The finding that several of the potential imprinted genes impact developmental processes, lends additional support for the robustness of my approach. I also covered examples of how I could deduce the phenotypes of the potential imprinted genes discovered by my approach. Nonetheless, only experimental validations could demonstrate the strength of my approach. Therefore, I offer links for accessing and downloading my data on the positions of ZFBS and ZFBS-Morph overlaps [13], peaks in the density-plots [15], and the MLL1 morphemes in the build hg19 of the human genome [63].

(Supplemental figures are after the reference)

Supplemental Figures
(Additional references are listed at the end of this file)

**Fig. S1.**
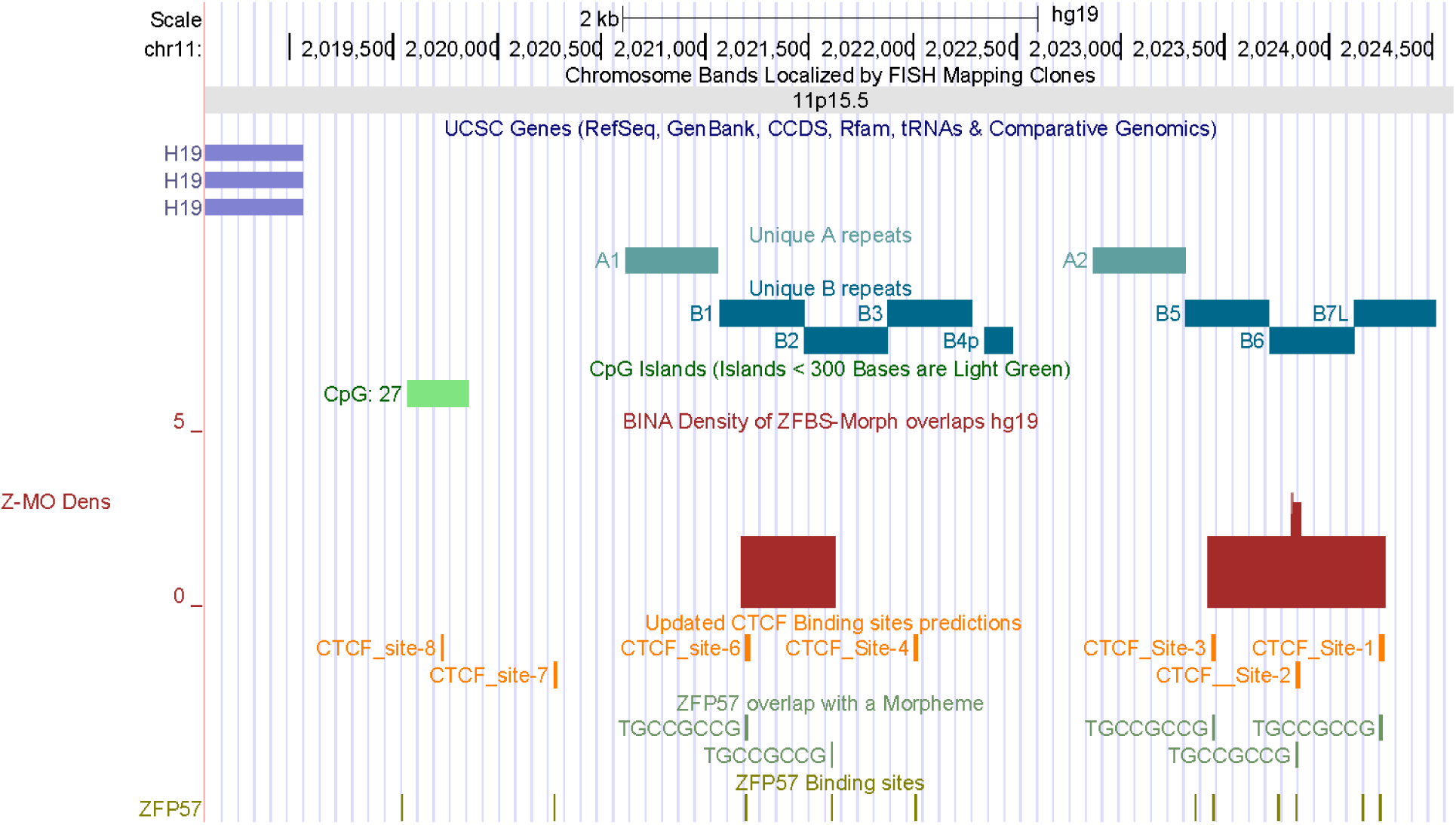
The position of a density peak consisting of a doublet located the ICR of *H19* – *IGF2* imprinted domain. See reference [1] for details about the positions of the unique repeats and CTCF sites upstream of the *H19* gene.

**Fig. S2.**
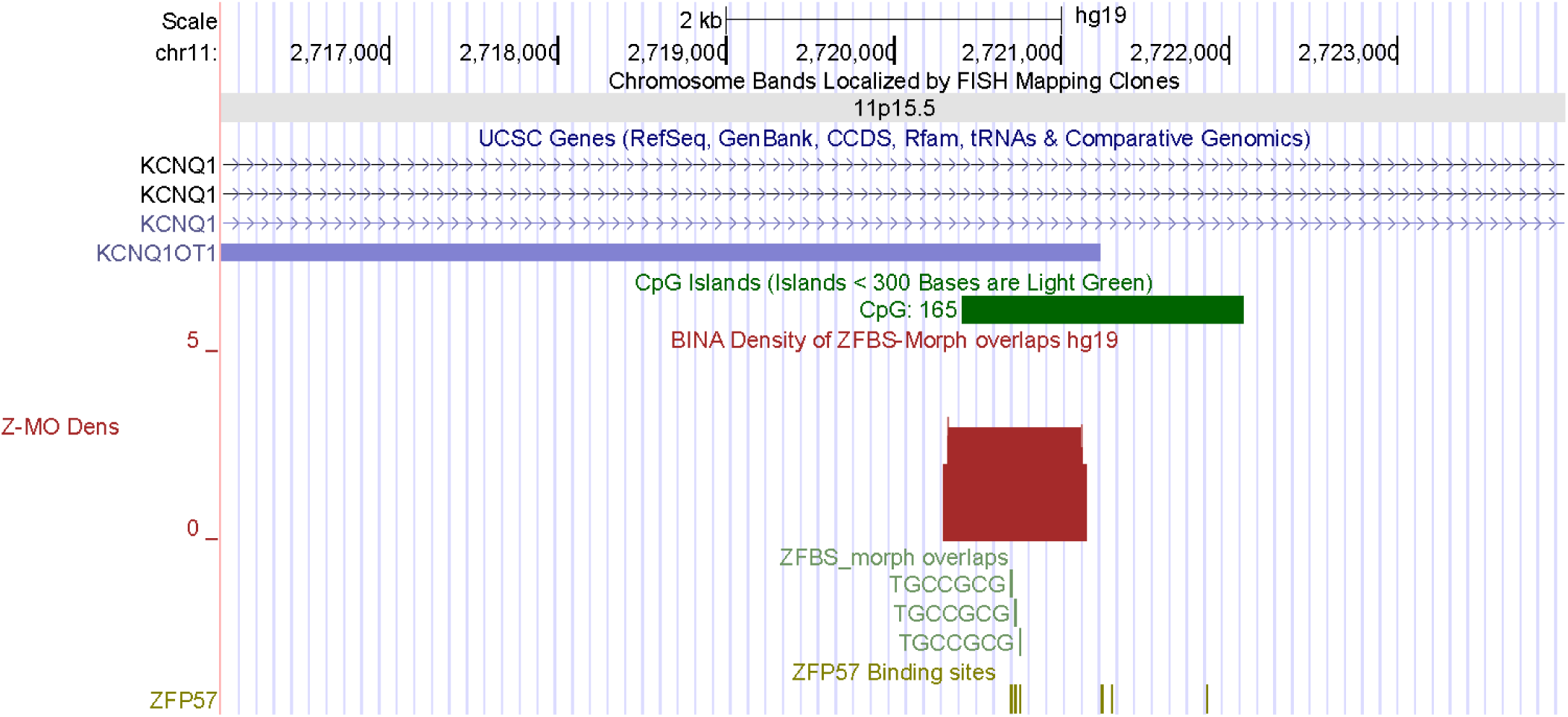
A peak in the density plots located the KvDMR in the *KCNQ1* imprinted domain.

**Fig. S3.**
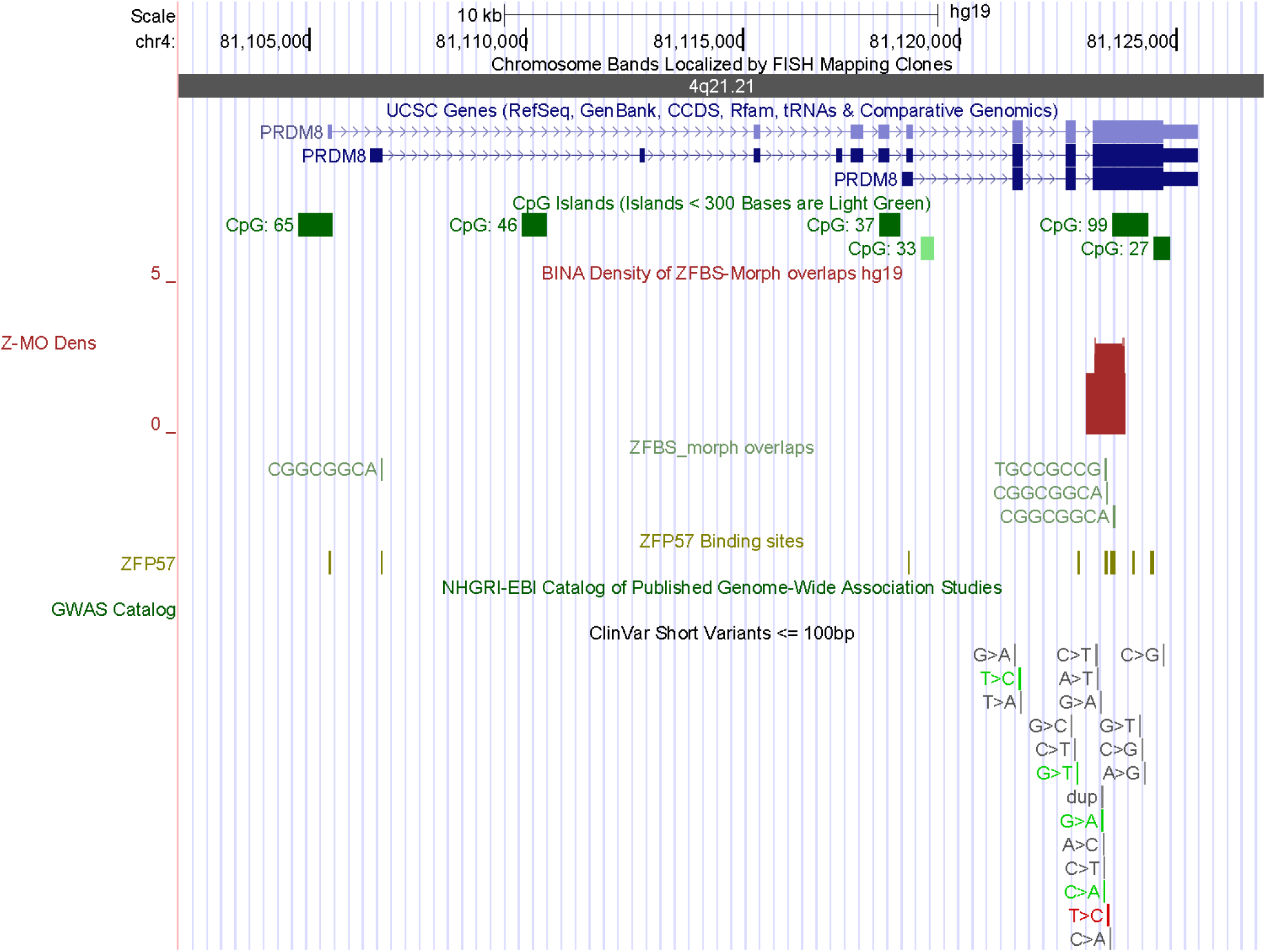
A peak in the density plots predicted an ICR for regulating the expression of a candidate imprinted gene deduced by experimental techniques [2]. This gene (*PRDM8*) is associated with neurodegenerative disorder characterized by onset of progressive myoclonus, ataxia, spasticity, dysarthria, and cognitive decline in the first decade of life (https://www.omim.org/entry/616640).

**Fig. S4.**
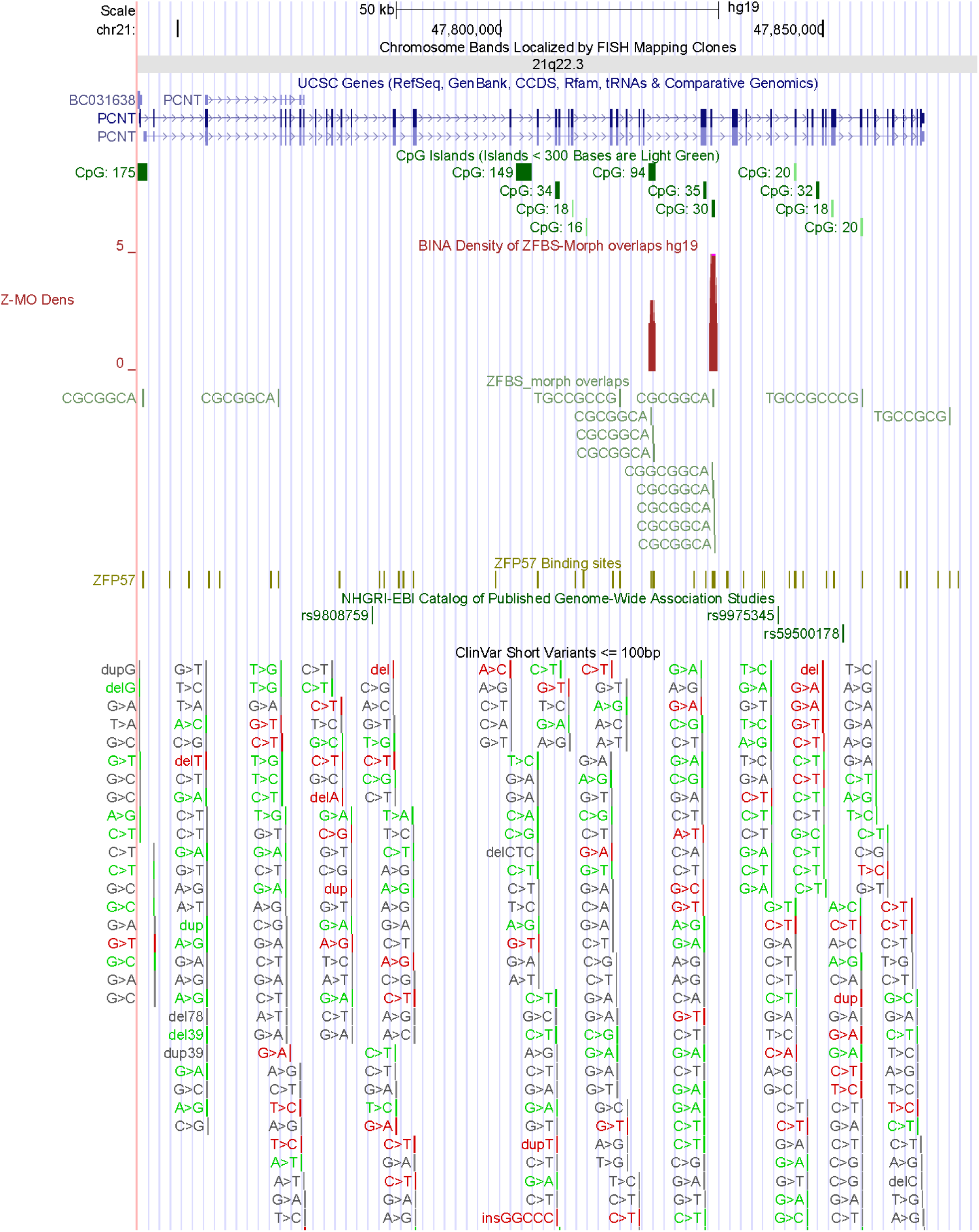
In the density plots,a peak predicted an ICR for regulating the expression of a candidate imprinted gene deduced by experimental techniques [2]. Homozygous or compound heterozygous mutations in *PCNT* caused Microcephalic osteodysplastic primordial dwarfism type II (https://www.omim.org/entry/210720).

**Fig. S5.**
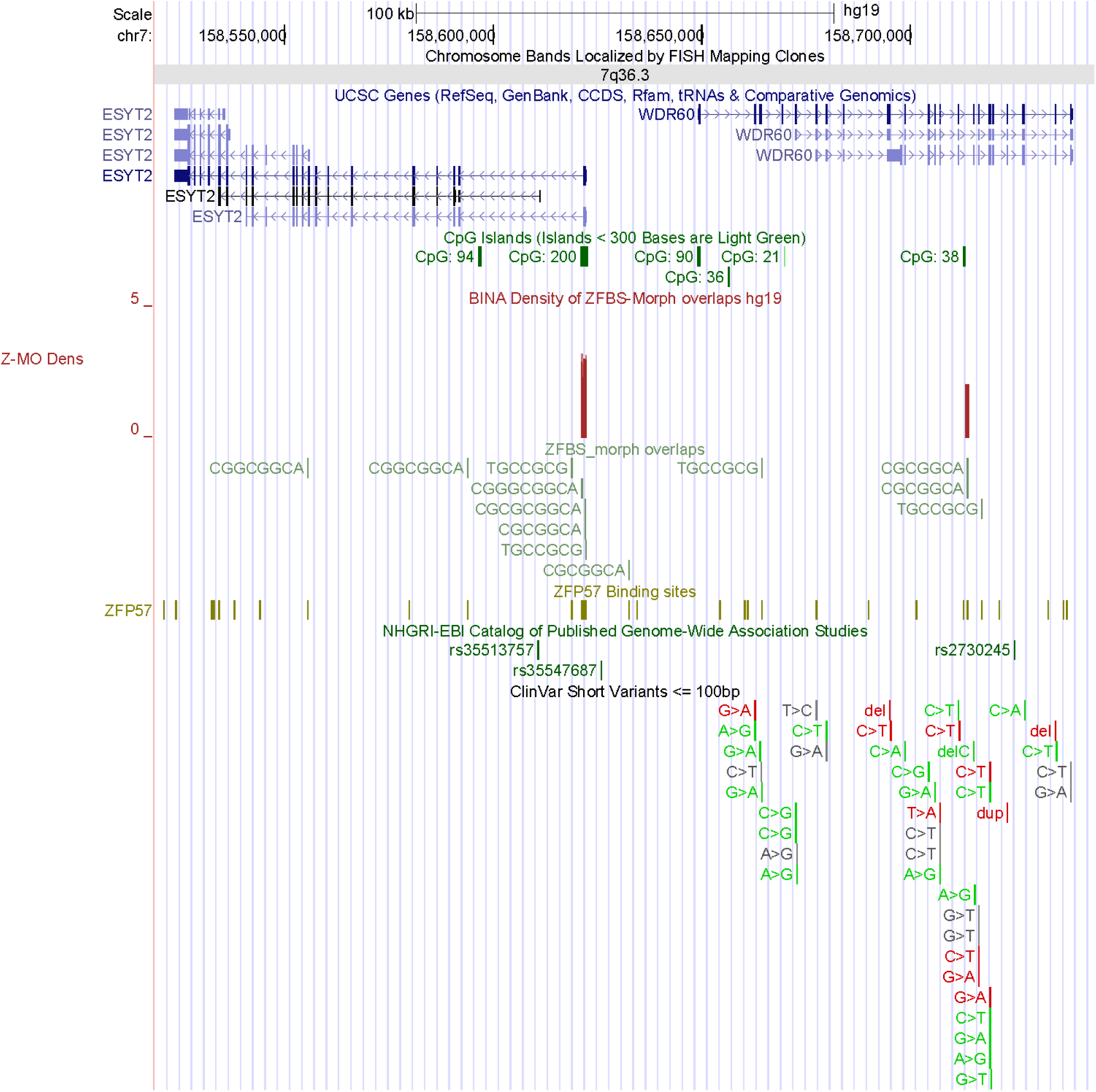
In the density plots, a peak predicted an ICR for regulating the expression of an imprinted gene deduced by experimental techniques [2]. This gene (*WDR60*) is associated with Short-rib thoracic dysplasia 8 (SRTD8) with or without polydactyly (https://www.omim.org/entry/615503). Note that the peak within *WDR60* covers 2 ZFBS-morph overlaps. Therefore, this peak could be a true or a false-positive. If false-positive, then another peak -far upstream of (*WDR60*)-could be a candidate ICR for regulating parent-of-origin specific expression of both *WDR60* and *ESYT2*. The latter gene encodes a protein (synaptotagmin-like protein 2) that belongs to a family of membranous Ca2^+^-sensors [3].

**Fig. S6.**
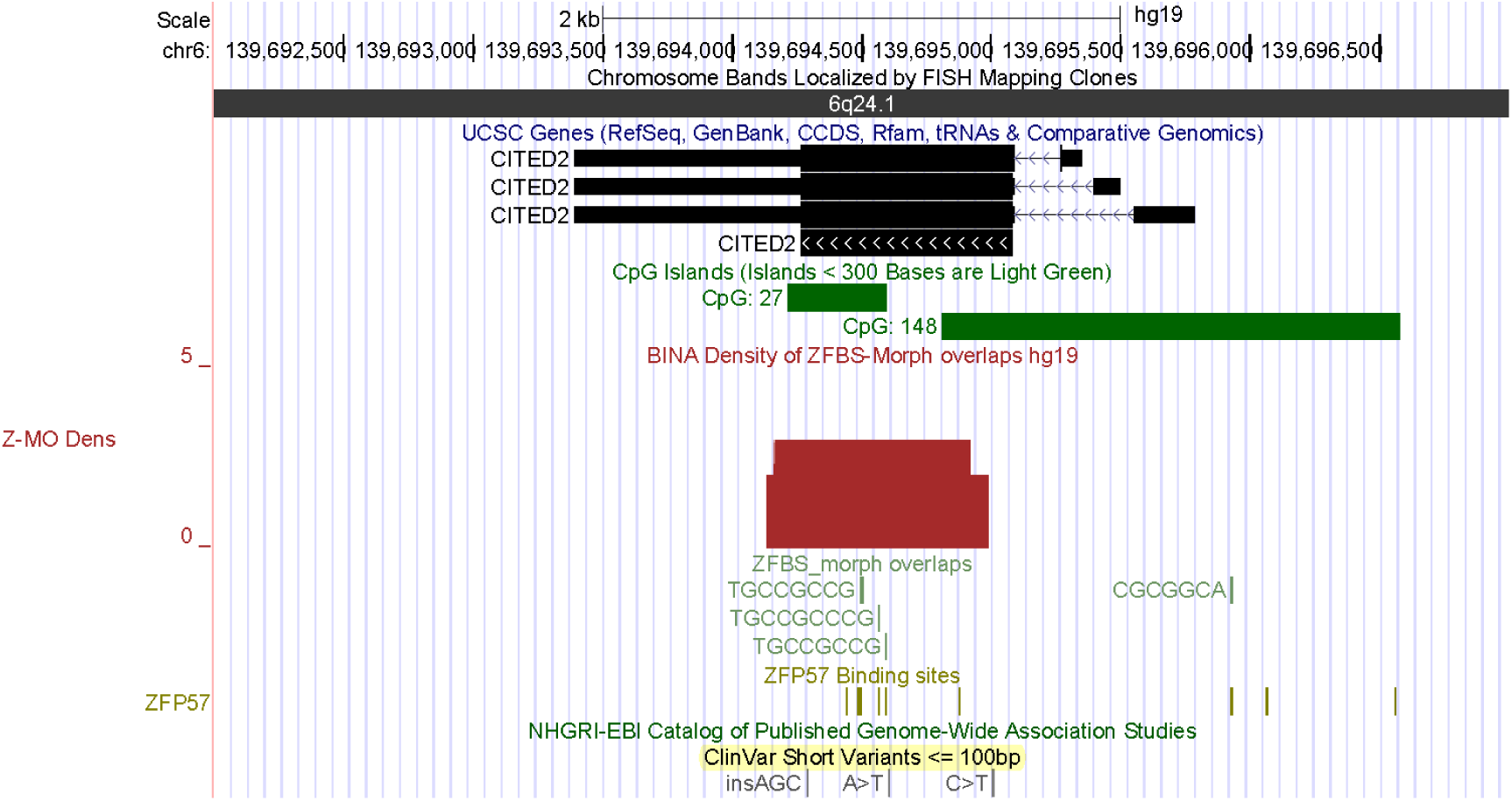
A density peak predicted a candidate ICR for a potential imprinted gene (*CITED2*). This gene encodes a regulator of transcription. Absence of *Cited2* in mouse embryos caused congenital heart disease by perturbing left-right patterning of the body axis [4]. Deleterious mutations in the human gene (*CITED2*) cause ventricular septal defect 2 (VSD2). This disorder is the most common form of congenital cardiovascular anomaly. It occurs in nearly 50% of all infants with a congenital heart defect and accounts for 14% to 16% of cardiac defects that require invasive treatment within the first year of life (https://www.omim.org/entry/614431).

**Fig. S7.**
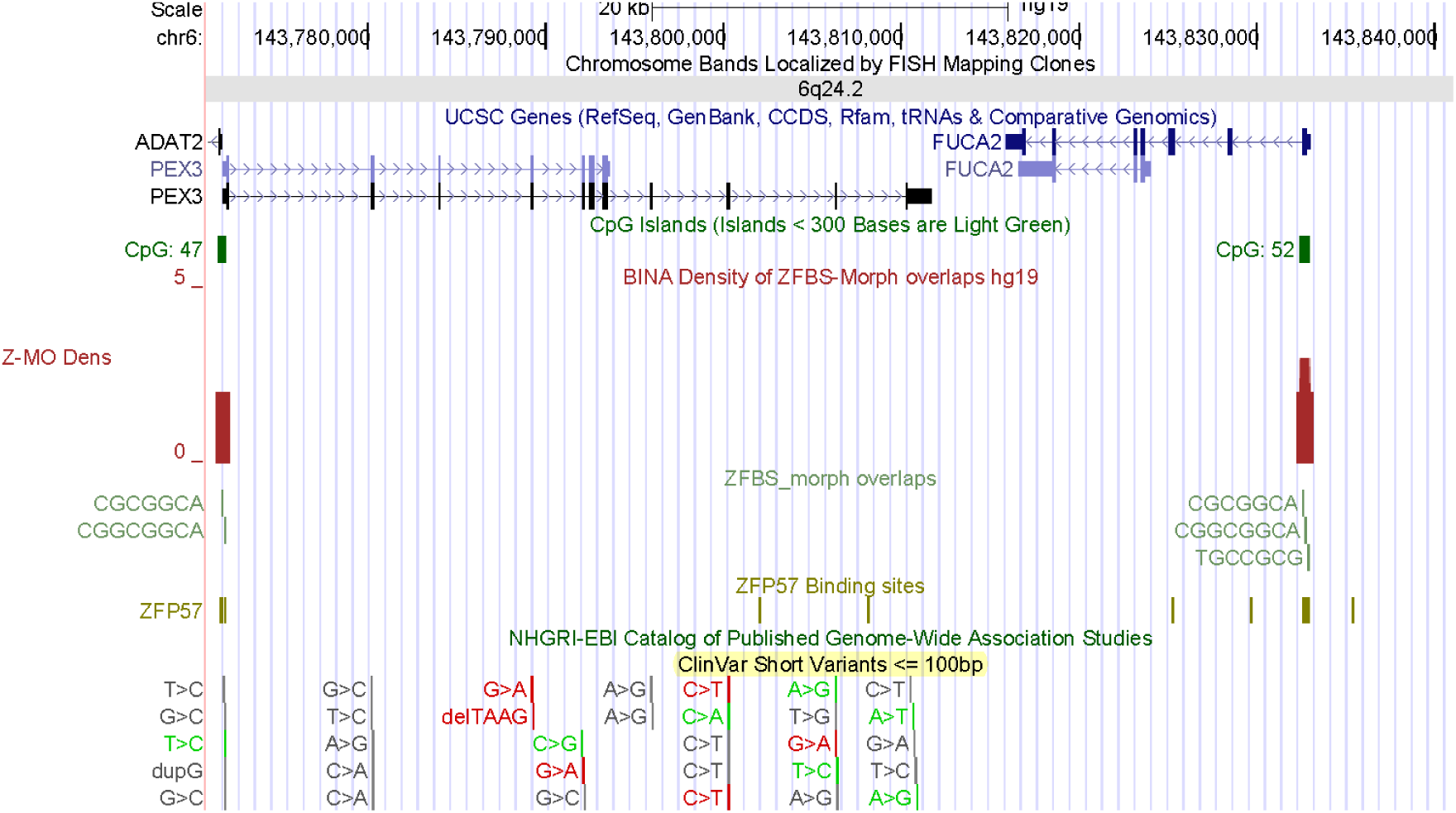
A density peak predicted a candidate ICR for a potential imprinted gene (*FUCA2*). Genome-wide analyses have identified *FUCA2* and *IL-18* as novel genes associated with diastolic function in African Americans with sickle cell disease [5].

**Fig. S8.**
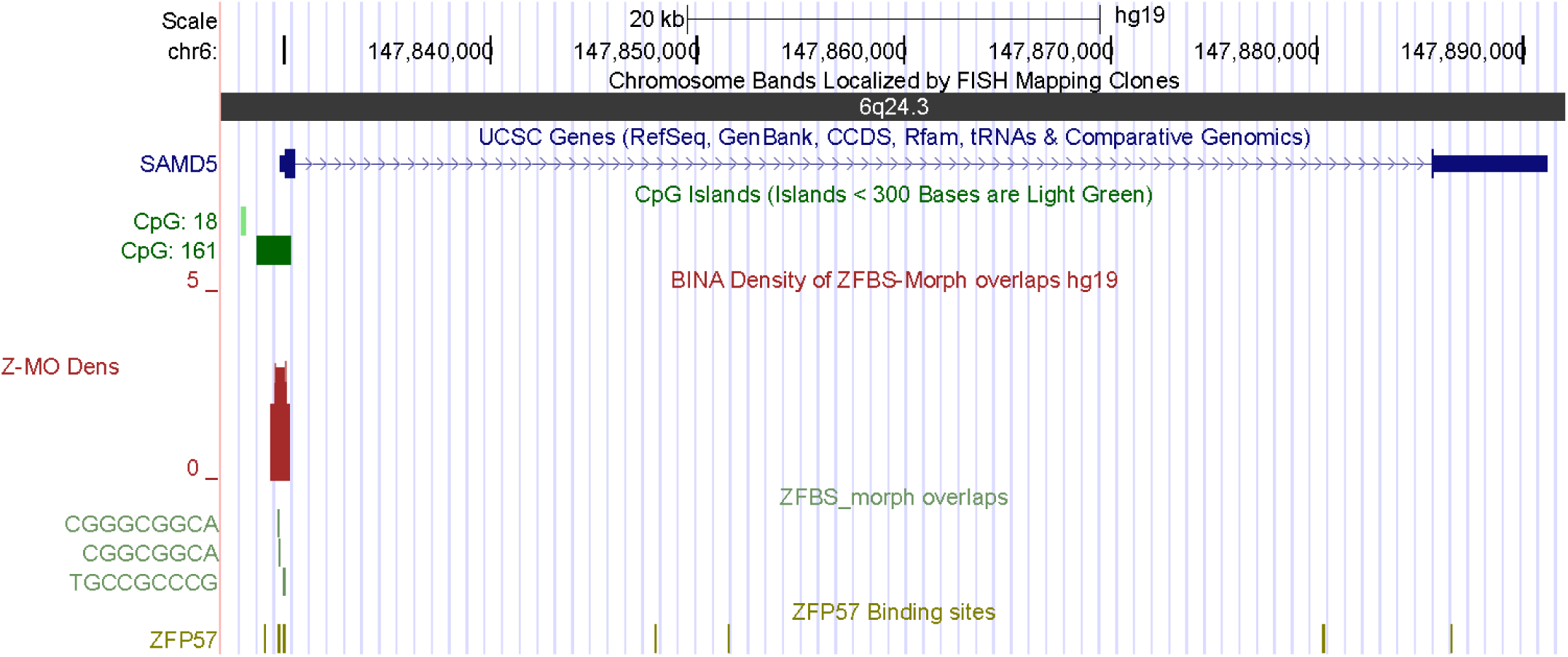
A density peak predicted a candidate ICR for a potential imprinted gene (*SAMD5*). A study found that *SAMD5* was overexpressed in prostate cancer and had powerful prognostic ability for predicting post-operative biochemical recurrence after radical prostatectomy [6].

**Fig. S9.**
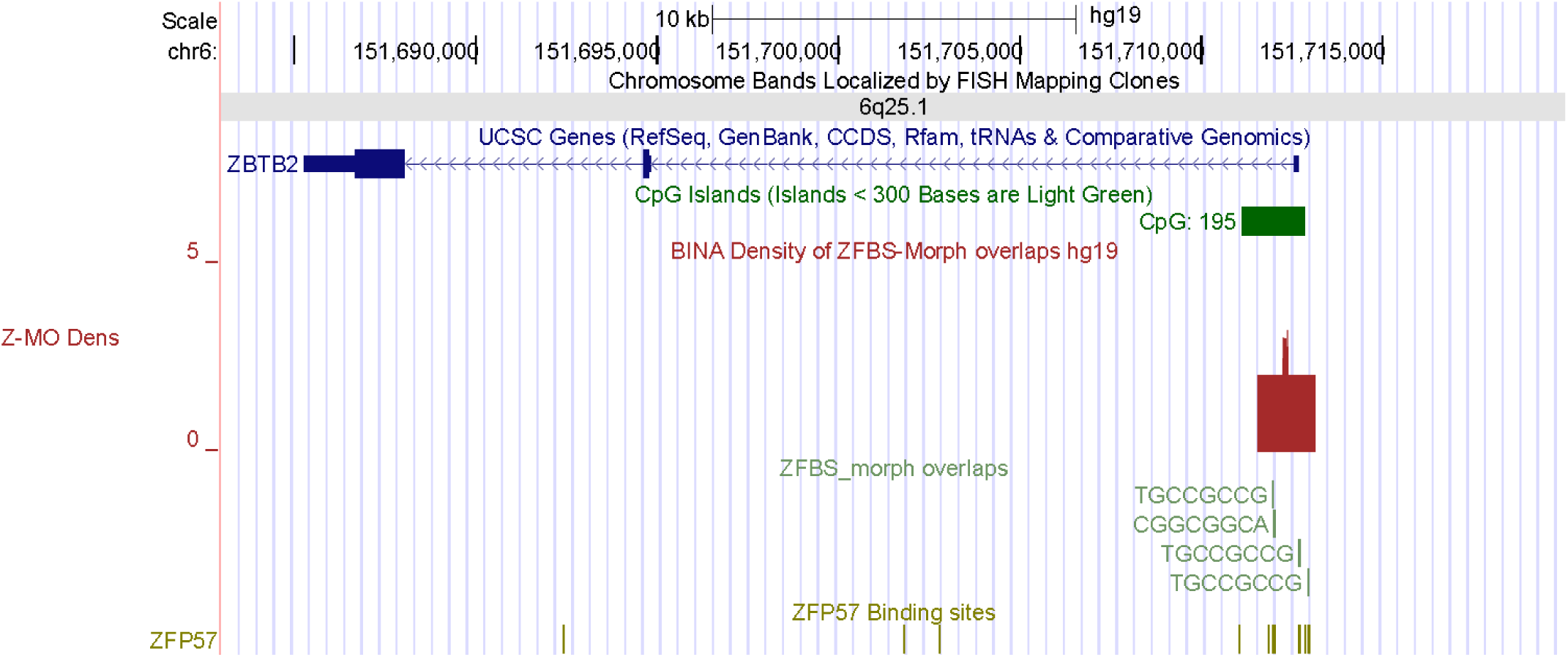
A density peak predicted a candidate ICR for a potential imprinted gene (*ZBTB2*). In mouse embryonic stem cells, ZBTB2 dynamically interacted with nonmethylated CpG island promoters and regulated differentiation [7]. In colorectal cancer, the abnormal forms of *ZBTB2* increased cell proliferation [8].

**Fig. S10.**
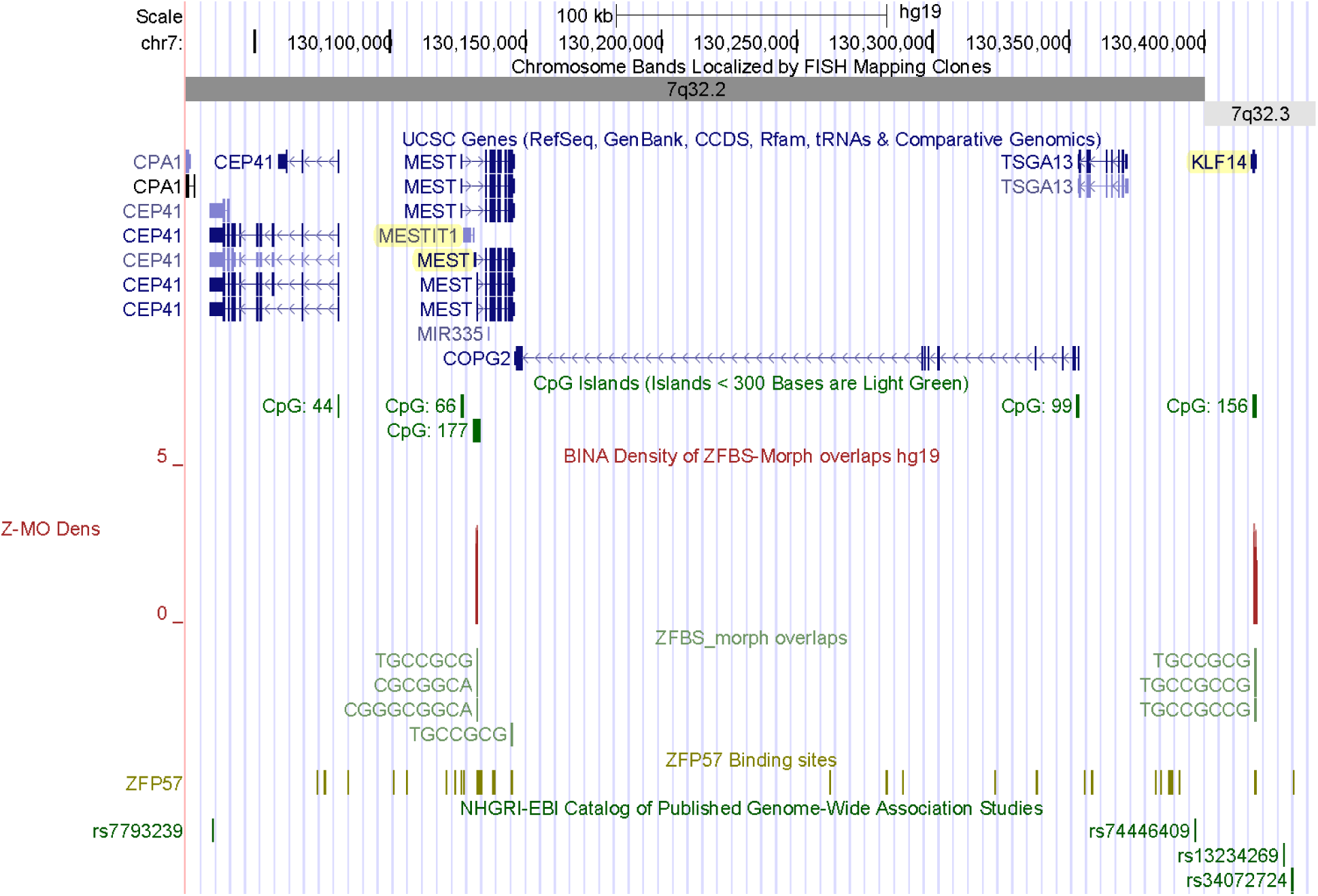
Density peaks mapping to known parent-of-origin specific transcripts. The density peak correctly located the ICR in the *MEST* locus. This ICR is intragenic and encompasses the TSSs of *MESTIT1* (a noncoding RNA gene) and a subset of *MEST* transcripts. *KLF14* is a known imprinted gene [9]. Density-plots predicted a candidate ICR regulating its imprinted expression.

**Fig. S11.**
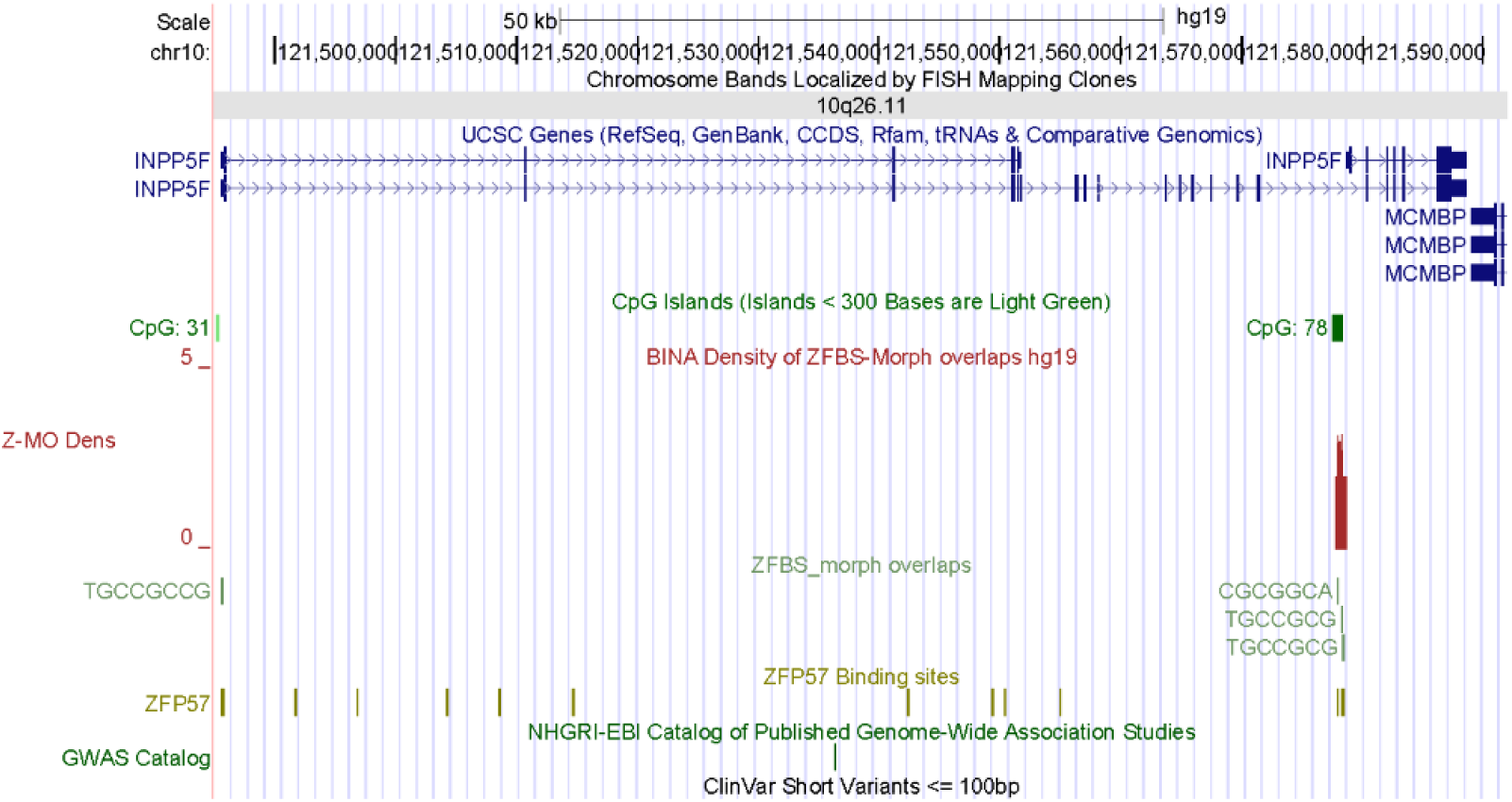
In density-plots, a peak correctly located the ICR regulating the expression of *INPP5_v2*.

